# Detection of *Mycobacterium tuberculosis* in human tissue via RNA *in situ* hybridization

**DOI:** 10.1101/2023.10.04.560963

**Authors:** Kievershen Nargan, Threnesan Naidoo, Mpumelelo Msimang, Sajid Nadeem, Gordon Wells, Robert L Hunter, Anneka Hutton, Kapongo Lumamba, Joel N Glasgow, Paul V Benson, Adrie JC Steyn

## Abstract

**Rationale:** Accurate TB diagnosis is hampered by the variable efficacy of the widely-used Ziehl-Neelsen (ZN) staining method to identify *Mycobacterium tuberculosis* (*Mtb*) acid-fast bacilli (AFB). Here, we sought to circumvent this current limitation through direct detection of *Mtb* mRNA.

**Objectives:** To employ RNAscope to determine the spatial distribution of *Mtb* mRNA within tuberculous human tissue, to appraise ZN-negative tissue from confirmed TB patients, and to provide proof-of-concept of RNAscope as a platform to inform TB diagnosis and *Mtb* biology.

**Methods:** We examined ante- and postmortem human TB tissue using RNAscope to detect *Mtb* mRNA and a dual ZN/immunohistochemistry staining approach to identify AFB and bacilli producing antigen 85B (Ag85B).

**Measurements and main results:** We adapted RNAscope for *Mtb* and identified intact and disintegrated *Mtb* bacilli and intra- and extracellular *Mtb* mRNA. *Mtb* mRNA was distributed zonally within necrotic and non-necrotic granulomas. We also found *Mtb* mRNA within, and adjacent to, necrotic granulomas in ZN-negative lung tissue and in Ag85B-positive bronchial epithelium. Intriguingly, we observed accumulation of *Mtb* mRNA and Ag85B in the cytoplasm of host cells. Notably, many AFB were negative for Ag85B staining. *Mtb* mRNA was observed in ZN-negative antemortem lymph node biopsies.

**Conclusions:** RNAscope has diagnostic potential and can guide therapeutic intervention as it detects *Mtb* mRNA and morphology in ZN-negative tissues from TB patients, and *Mtb* mRNA in ZN-negative antemortem biopsies, respectively. Lastly, our data provide evidence that at least two phenotypically distinct populations of *Mtb* bacilli exist *in vivo*.

## INTRODUCTION

Tuberculosis (TB) continues to be a threat to global health, with significant morbidity and mortality. Thus, accurate and rapid detection of *Mtb* bacilli is critical for definitive TB diagnosis and research efforts. Culturing of isolates remains the gold standard for TB diagnosis but takes weeks to complete. Instead, detection of *Mtb* in clinical samples has traditionally relied on the microscopic observation of acid-fast bacilli (AFB) following Ziehl-Neelsen (ZN) staining. However, since *Mtb* can be cultured from ZN-negative tissue specimens, ZN staining can produce false negatives, delaying therapeutic intervention (1, 2). Further, ZN staining provides no insight into the physiological state of *Mtb, i.e*., whether the bacillus is alive or dead. Importantly, studies in the 1940s showed that single *Mtb* colonies are stratified into three layers; nonacid fast, weakly acid-fast, and strongly acid-fast bacilli (3). Also, in the 1950s, Canetti reported that bacillary destruction generates “ghosts” of non-AFB in the caseum, which became scarcer as the age of necrosis increased (4). Unfortunately, the implications of this important biological property, i.e., acid-fastness, have been underappreciated. Since *Mtb* can transition from ZN-positive to ZN-negative *in vivo* and in response to drug therapy (5), there is a strong unmet need for improved detection of *Mtb*.

*In vitro* and *in vivo* studies have shown that several secreted *Mtb* antigens are present within TB lesions (6). Despite the potential of using antibodies to detect *Mtb* surface or secreted antigens (7–11), little progress has been made to improve TB diagnostics beyond ZN staining. Whereas mRNA-based diagnostic tests have been proposed for TB (12), studies comparing RNA *in situ* hybridization (RISH) methods with routine histochemistry and/or immunohistochemistry (IHC) for the spatial identification of *Mtb* are lacking, especially in paucibacillary and abacillary tissues. A novel RISH platform, referred to as RNAscope, was recently developed (13). RNAscope uses an innovative probe design to increase specificity and the signal-to-noise ratio and allows visualization of single mRNA molecules (as punctate dots) within cells in formalin-fixed paraffin-embedded (FFPE) tissue (13–17). Since the quantity of a specific *Mtb* mRNA molecule can exceed that of its corresponding gene by orders of magnitude, RNAscope may allow assessment of the spatial and microenvironmental distribution of mRNA *in vivo*. This knowledge may provide further insight into mycobacterial physiology and heterogeneity *in vivo*. Therefore, the goal of this study was to adapt RNAscope to detect *Mtb* in ZN-positive human pulmonary and extrapulmonary TB tissue and in ZN-negative ante- and postmortem tissue with proven active TB. Finally, we employed IHC to detect *Mtb* antigens to investigate whether *Mtb* bacilli in human tissue exist as a homogenous population regarding antigen production.

## MATERIALS AND METHODS

### Ethics and Human Subjects

This study was approved by the University of KwaZulu-Natal Biomedical Research Ethics Committee (BREC; class approval numbers BCA 535/16, and BE019/13) and the University of Alabama Birmingham Institutional Review Board (IRB; study numbers IRB-300008174 and IRB-300008174-2). Consent for the use of autopsy material for research is included in the UAB authorization for autopsy consent form. Patients undergoing lung resection for TB were recruited from King DinuZulu Hospital Complex, a tertiary center for patients with TB in Durban, South Africa. See Supplemental Materials and Methods and Table E1 in the online data supplement for description of human subjects and tissues.

### Histology and Immunohistochemistry

Human tissue specimens were aseptically removed and fixed in 10% neutral buffered formalin (10% NBF). Specimens were processed using a xylene-free protocol and embedded in paraffin wax. Tissue sections were cut and mounted on charged slides. Histological and immunohistochemical staining was performed essentially as described (18). Slides with stained tissues were imaged using a Hamamatsu NDP slide scanner (NanoZoomer RS2, Model C10730-12) or an Olympus BX50 microscope with a DP23 digital microscope camera. See Supplemental Materials and Methods in the online data supplement for additional details.

### RNAscope

The RNAscope 2.5 High Definition (HD)-Red assay kit (Advanced Cell Diagnostics (ACD) cat # 322350) uses a red chromogen to visualize target mRNA. We used a positive control RNAscope probe set of 16 probe pairs directed against bp 139-989 of human peptidylprolyl isomerase B (*PPIB*) mRNA (ACD cat # 313901). The negative control probe set consists of 10 probe pairs directed against bp 414-862 of *Bacillus subtilis* dihydrodipicolinate reductase (*dapB*) mRNA (ACD cat # 310043). The *Mtb*-specific probe set (ACD cat # 552911) consists of 120 probe pairs directed against mRNAs from six genes: secreted L-alanine dehydrogenase (*ald*), catalase-peroxidase-peroxynitritase T (*katG*), trehalose-6-phosphate phosphatase (*otsB1*), resuscitation-promoting factor A (*rpfA*), resuscitation-promoting factor B (*rpfB*), and ATP-binding protein (ABC transporter) (*irtB*). See Supplemental Materials and Methods in the online data supplement for additional details.

### HALO® analysis

Whole-slide image analysis was performed using Halogen-Assisted Light Optimization (HALO) v3.6.4134 (Indica Labs, Corrales, NM). The ISH module v4.2.3 (Indica Labs, Corrales, NM) was used to detect RNAscope signals and the Area Quantification module v2.4 was used for visualization. See Supplemental Materials and Methods in the online data supplement for additional details.

## RESULTS

### RNAscope cellular and tissue controls

RNAscope is a highly specific and sensitive RISH technique that combines multiple paired “Z” oligonucleotide probes with signal amplification steps to visually detect single mRNA molecules in tissue samples while preserving tissue architecture (13) (Figure 1A). We posited that RNAscope could detect mRNA in bacilli, infected host cells, and antemortem/postmortem human specimens (Figure 1B). RNAscope assays were initially validated by exposing thin sections of a Hela cell pellet to a human mRNA positive control probe set directed against peptidylprolyl isomerase B (*PPIB*) mRNA (Figure E1A) and a negative control probe set specific for *Bacillus subtilis* dihydrodipicolinate reductase (*dapB*) mRNA (Figure E1B) resulting in robust signals and no signal, respectively. The *Mtb*-specific RNAscope probe set (120 pairs) targets six mRNAs to provide improved specificity and sensitivity. We tested the *Mtb* probe set against human neonatal lung tissue, which showed no signal as expected (Figure E1C). Lastly, we tested the *PPIB* control probe against testicular tissue from a TB patient (Figure E2) to verify the integrity of human specimens for RNAscope analysis. Overall, these control assays demonstrate that RNAscope is compatible with the tissue processing protocols for our human specimens.

**Figure 1.**
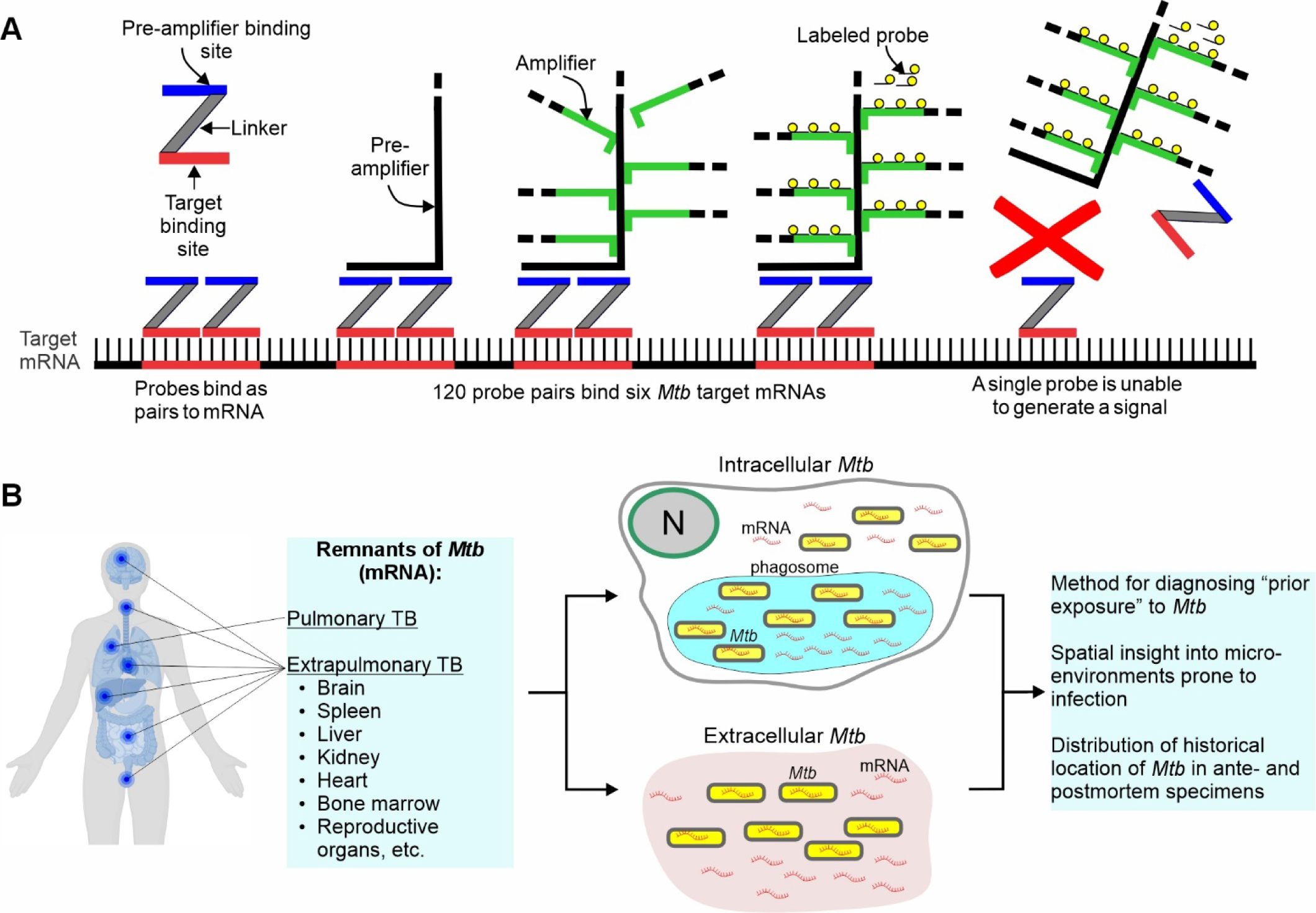
RNAscope and its application for detecting *Mtb* in human tissues. (**A**) Schematic of the RNAscope *in situ* hybridization platform for mRNA detection. Following tissue fixation and permeabilization, up to 20 paired “Z” oligonucleotide probes hybridize with multiple target sequences (red line) within a single mRNA across a ∼1,000-bp region. For detection of *Mtb* in human tissue, 120 paired probes bind six *Mtb* mRNAs. Subsequent steps include binding of a pre-amplifier to mRNA-bound paired “Z” probes, addition of multiple amplifiers molecules, and binding of paired probe-specific labels (fluorescence or chromogen based) for detection of mRNA via microscopy where an individual mRNA molecule appears as a punctate dot. Binding of a single “Z” probe does not result in a signal. (**B**) Diagram illustrating the potential of RNAscope to identify and track the historical footprints of remnants (mRNA) of *Mtb* in pulmonary, extrapulmonary, and postmortem tissue. Schematic in **(A)** adapted from (40).

### RNAscope detects intact and disintegrated *Mtb* bacilli, and mRNA in human tissue

To compare the ability of RNAscope and ZN methods to detect *Mtb* bacilli *in situ*, we examined seminiferous tubules containing AFB obtained from a patient with active TB. Since the structurally complex *Mtb* cell wall is a barrier (5) to RNAscope probes, conditions were optimized to allow *Mtb*-directed RNAscope probes to enter intact bacilli. RNAscope identified *Mtb* bacillary rods inside and outside the seminiferous tubules that are visually similar to ZN-stained bacillary rods (Figure 2A-F). RNAscope revealed a spectrum of signal shapes: bacilli with halos, bacillus-sized dots (consistent with the diameter of a vertical bacillus within the cut plane), diffused halos, and smaller punctate dots (Figure 2F, G, H-M), which were also observed within host cells (Figure 2G). RNAscope signal intensity (faint to prominent) is proportional to the number of paired probes bound to target mRNAs, not transcript abundance.

**Figure 2.**
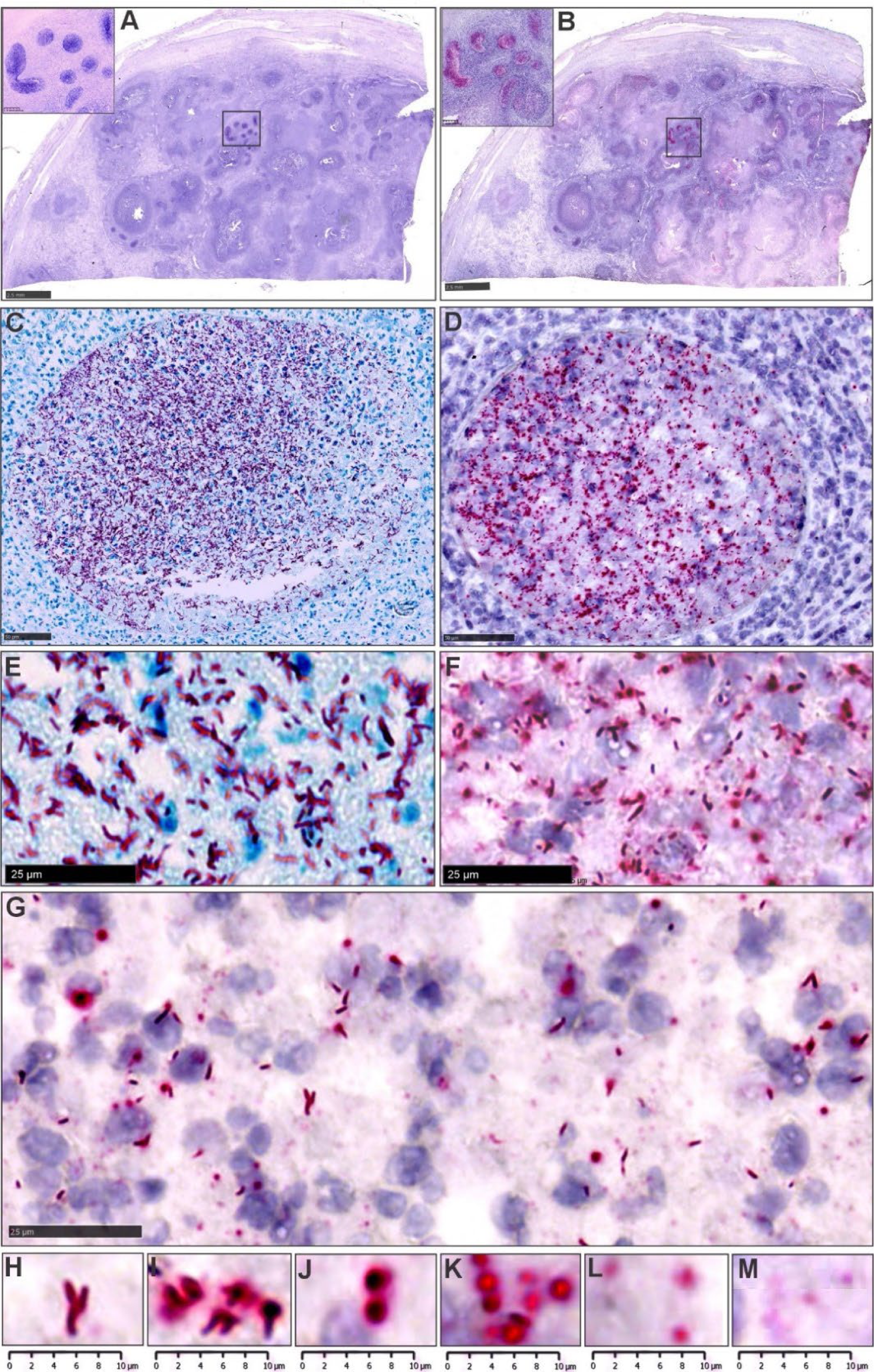
RNAscope can detect intact and disintegrated *Mtb* bacilli and mRNA in human tissue. (**A**) low power image of a human testicular specimen revealing ZN-positive AFB. Inset - note positive ZN staining within the seminiferous tubules. (**B**) low power image of the same human testicular specimen revealing *Mtb*-specific RNAscope signals, which are abundant within the seminiferous tubules. Medium power image revealing ZN-positive AFB (**C**) and *Mtb*-specific RNAscope signal (**D**) within seminiferous tubules. High power image revealing ZN-positive AFB (**E**) and *Mtb*-specific RNAscope signal (**F**). (**G**) High power image showing a spectrum of *Mtb* RNAscope signals within a single field **(H-M)** High power images of RNAscope signals from (**H**) intact AFB, (**I, J**) partially intact bacilli, (**K, L**), large, diffuse dots containing concentrated *Mtb* RNA from disintegrated bacilli. (**M**) small dots representing single *Mtb* mRNA transcripts.

In sum, improved permeabilization permits RNAscope probes to detect *Mtb* bacillary rods. Secondly, a halo surrounding bacilli suggests RNA leakage, indicating cellular disintegration that supports Canetti’s findings of bacillary disintegration and loss of acid-fastness *in vivo* (4). Thirdly, our findings show that *Mtb* mRNA is stable enough for detection with RNAscope in numerous microenvironments. These findings provide new biological insight into bacillary morphology *in vivo* and cell death, which cannot be obtained through ZN staining.

### RNAscope detects *Mtb* mRNA in ZN-negative human TB lung tissue

We next examined the utility of RNAscope for identifying *Mtb* or remnants of infection, *i.e*., mRNA, in ZN-negative human TB tissue. ZN staining of formalin-fixed, paraffin-embedded (FFPE) lung tissue from a patient with active TB showed numerous extracellular and intracellular bacilli (Figure 3A). However, a different FFPE lung tissue specimen from the same patient was ZN-negative. This unsurprising as most human pulmonary TB granulomas contain few, if any, ZN-positive *Mtb*. In contrast, RNAscope analysis of this ZN-negative tissue revealed numerous RNAscope signals within necrotic granulomas (Figure 3B, C). RNAscope signals were also abundant in bronchial epithelial cells (Figure 3D) which were ZN-negative (Figure 3E) and antigen 85 (Ag85B)-positive (Figure 3D).

**Figure 3.**
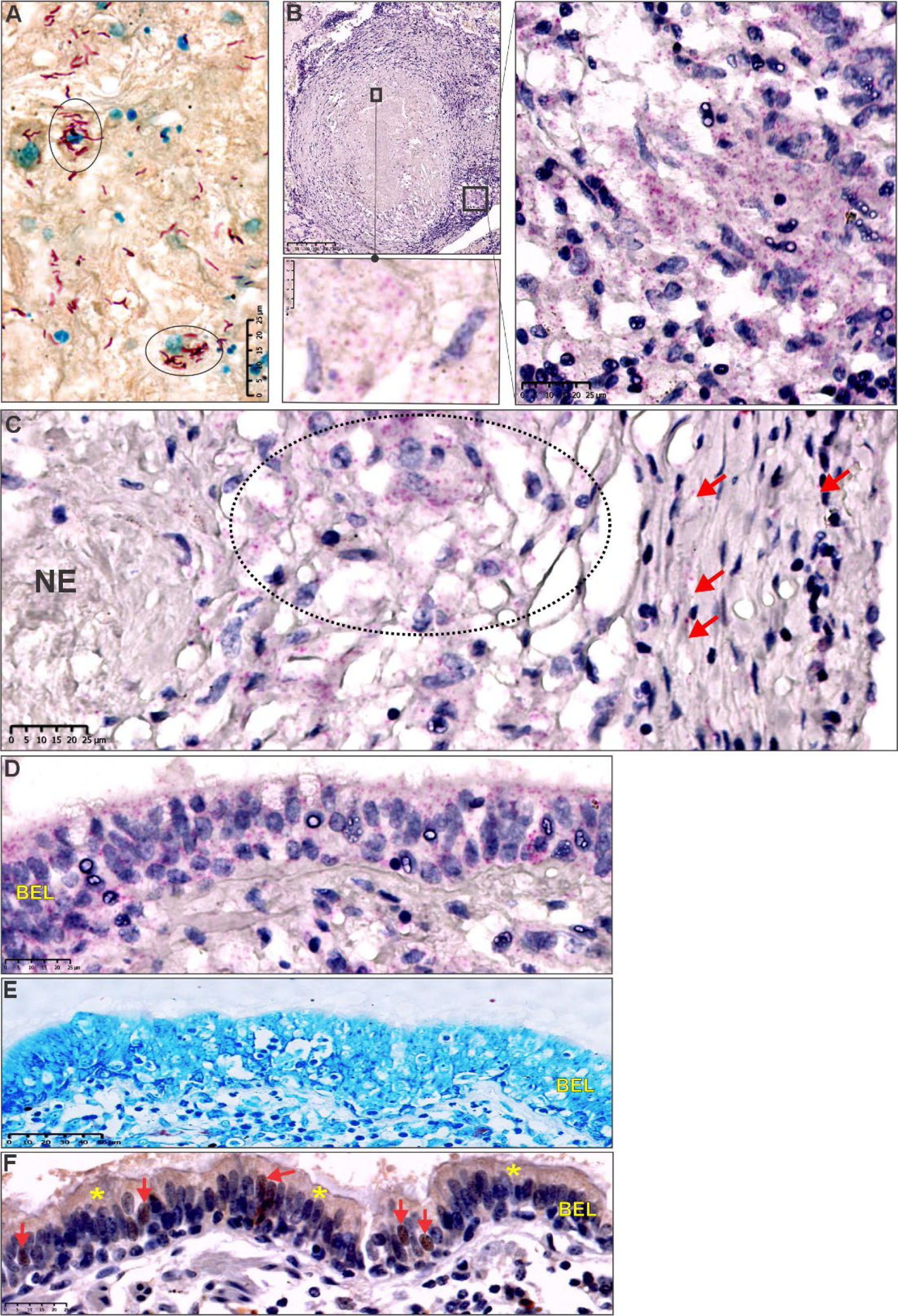
RNAscope detects *Mtb* mRNA in human TB lung tissue. Lung tissue from a patient with active TB. (**A**) ZN-positive tissue section revealing numerous intracellular (circled) and extracellular *Mtb*, (**B**) ZN-negative tissue containing a necrotic granuloma with RNAscope signal positive for *Mtb* mRNA: Boxed regions correspond to high power images showing *Mtb* mRNA. (**C**) High power image revealing RNAscope signals within a necrotic area (NE), granulomatous area (oval) and granulomatous inflammatory layer (red arrows). (**D**) high power image of numerous RNAscope signals within the bronchiolar epithelial layer (BEL). (**E**) ZN stain showing the absence of AFB. (**F**) High power image showing Ag85B-positive staining within the cytoplasm (yellow asterisks) and nuclei (red arrows) of bronchiolar epithelial cells.

These data demonstrate that *Mtb* mRNA is abundantly present in ZN-negative lung tissue sections of a confirmed case of pulmonary TB and that *Mtb* mRNA is sufficiently stable to be detected by RNAscope. RNAscope and Ag85B immunostaining suggest the bronchial epithelial layer as an overlooked entry portal for *Mtb* infection.

### *Mtb* mRNA and secreted antigens accumulate in the cytoplasm of host cells

In a study of individuals who died from causes other than TB, *Mtb* DNA was found in alveolar and interstitial macrophages, type II pneumocytes, endothelial cells, and fibroblasts (19). While AFB were not clearly identified in these cells, these findings are consistent with early reports (20–23) that *Mtb* can persist in lung tissue without histological evidence of TB lesions. Consistent with those studies, we identified accumulated *Mtb* mRNA in the cytoplasm of several alveolar epithelial cells (Figure 4A-D) in a signal pattern distinct from the intracellular punctate dots that represent single *Mtb* mRNA molecules. We refer to such cells as <u>h</u>ost-cell <u>a</u>ccumulated *<u>Mt</u>b* <u>R</u>NA (HAMR) cells*. Mtb* mRNA is present in HAMR cells within adjacent (Figure 4B, C), consolidated alveolar areas (Figure 4D), and within lymphocytic aggregates at the periphery of necrotic granulomas (Figure 4E). The functional significance of *Mtb* mRNA within HAMR cells is unknown; however, these mRNAs may act as pathogen-associated molecular patterns (PAMPs) as has been shown for mRNAs in conventional innate immune cells (24–28) to modulate immunity.

**Figure 4.**
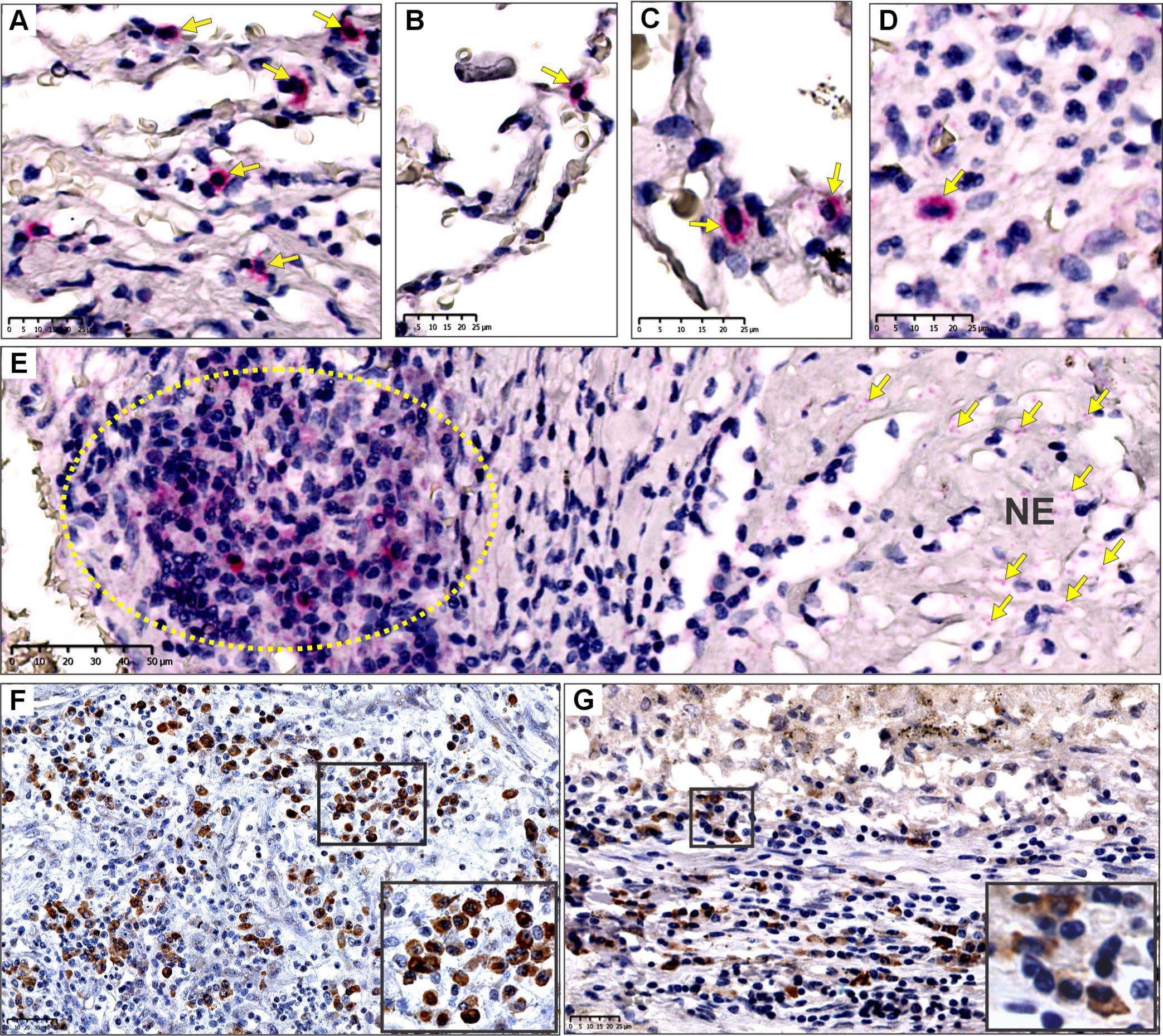
Accumulation of *Mtb* mRNA and secreted antigens in cells from pulmonary and extrapulmonary tissue. (**A-D**) high power images showing prominent RNAscope signals of *Mtb* mRNA within alveolar epithelial cells (yellow arrows). (**E**) high power image of RNAscope signals within lymphoid aggregates of necrotic granulomas (area in yellow oval). **(F)** Medium power image of Ag85B-positive staining in the cytoplasm of macrophages in extrapulmonary (testicular) human tissue specimens. (**G**) Medium power image of Ag85B-positive staining in the cytoplasm of alveolar macrophages and epithelioid histiocytes within pulmonary human TB specimens. Insets show higher power images of the cytoplasmic localization of Ag85B staining.

Secreted *Mtb* antigens have been identified via IHC staining in *Mtb*-infected human lung tissue (7–11). Similarly, we used IHC to detect *Mtb* secreted antigen 85B (Ag85B) in the same pulmonary and testicular specimens used for RNAscope analysis (Figures 2, 3). We observed Ag85B accumulation in numerous cell types (Figure 4F) including alveolar macrophages and epithelioid histiocytes (Figure 4G). In sum, our data demonstrate that cells from histologically normal and diseased airways accumulate *Mtb* mRNA and secreted antigens.

### Spatial distribution of *Mtb* RNA in host microenvironments

Our findings imply that RNAscope can identify *Mtb* mRNA as pathogenic “footprints” to reveal the history of infection, which is not possible with ZN staining. Since each punctate dot represents one *Mtb* mRNA molecule, accurate data interpretation requires quantitation. We used the HALO® image analysis platform to quantify *Mtb* mRNA signals in necrotic and non-necrotic granulomas, terminal and pulmonary bronchi, lymphocytic aggregates, and adjacent tissue from resected TB lung tissue (Figure 5A). To avoid potentially confounding variables such age, sex, tissue pathology bias, and experimental variability, we selected a single ZN-negative TB tissue specimen that contains all the aforementioned microenvironments.

**Figure 5.**
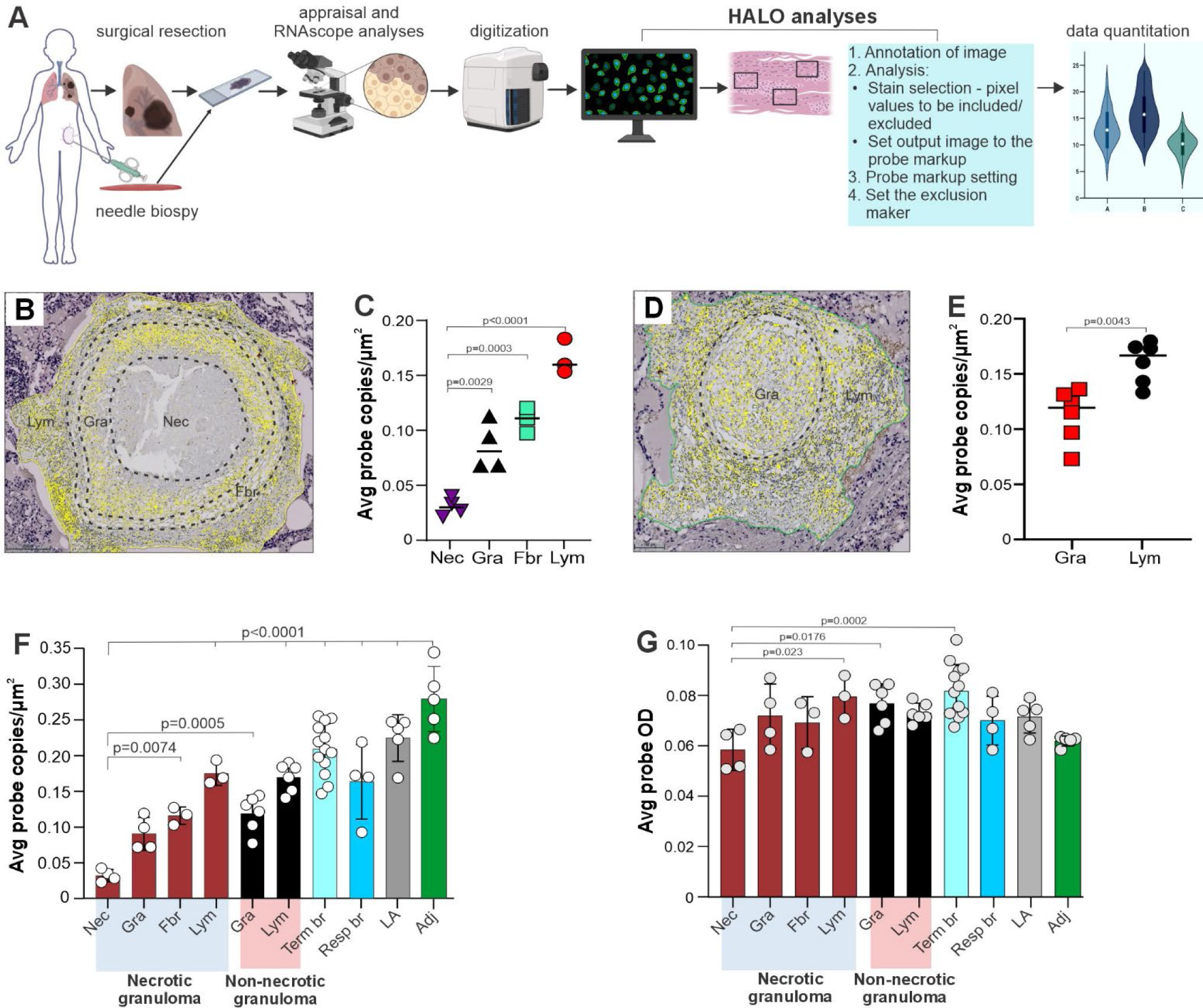
Quantification of RNAscope signals to inform the biology of TB lesions. (**A**) Flow diagram depicting sample acquisition, digitization, quantitation using HALO**®** software, and data interpretation. (**B**) Zonation of necrotic granulomas and (**C**) average number of RNAscope signals in each zone per square micrometer. (**D**) Zonation of non-necrotic granulomas and (**E**) average RNAscope signal optical density (OD) in each zone. (**B**, **D**) Yellow dots; RNAscope signals. (**F**) and (**G**) Average probe copies per square micrometer and average probe density (OD), respectively, in TB lesions and tissue. Zones include necrotic (Nec), granulomatous inflammatory (Gra), fibrotic (Fbr), lymphocytic (Lym), terminal bronchi (Term br), respiratory bronchi (Resp br), lymphocytic aggregates (LA) and tissue adjacent to diseased areas (Adj). Data in (**C**), (**E**), (**F**) and (**G**) represent the mean ± SD. In (**C**, **E**, **F and G**), each point represents the number of RNAscope signals per square micrometer (**C**, **E** and **F**) or probe OD (**G**) in an individual zone within a granuloma or other lung feature. In (**C, F, G**), four necrotic granulomas were examined that exhibited Nec and Gra zones (n=4) and Fbr and Lym zones (n=3). In (**E, F and G**), six non-necrotic granulomas were examined that contained Gra and Lym zones (n=6). (**F**, **G**) Term br; n=13, Resp br; n=4, LA; n=5, Adj; n=5. Data were analyzed using one-way ANOVA and Bonferroni’s multiple comparison test (**C**, **F** and **G**) or by unpaired Mann-Whitney test (**E**). (**C**, **F and G**) all comparisons are with respect to the Nec zone.

We hypothesized that quantifying RNAscope signals within four distinct regions of the necrotic granuloma (necrotic, granulomatous inflammatory, fibrotic, and lymphocytic; Figure 5B) could help inform the pathophysiology of granuloma formation. The necrotic zone had the lowest average probe copy (APC) per square micrometer, followed by progressive increases in APC within the granulomatous inflammatory, fibrotic, and lymphocytic zones (Figure 5C), suggesting that *Mtb* mRNA is distributed in accordance with discrete pathophysiological zones that may contribute to granuloma formation. We classified a non-necrotic granuloma (Figure 5D) into lymphocytic and non-necrotic zones and show that the APC in the lymphocytic zone is significantly higher than in the non-necrotic zone (Figure 5E).

*Mtb* mRNA signals were quantified in terminal and pulmonary bronchi, adjacent tissue, lymphocytic aggregates distant from necrotic or non-necrotic granulomas. Comparison of these APC values with those from necrotic and non-necrotic lesions indicates that the adjacent tissue, lymphocytic aggregates, and terminal bronchi exhibit the highest APC (Figure 5F). Terminal bronchi exhibited the highest average probe optical density (OD) (Figure 5G), indicating longer, more stable *Mtb* mRNAs that can bind more RNAscope probes.

In sum, the zonal distribution of *Mtb* mRNA within the granuloma offers new information into the evolution of granuloma formation and clinical course of disease that ZN staining cannot. Lastly, the spatial distribution of *Mtb* mRNA provides new insight into the historical footprints of *Mtb* within human tissue.

### Identification of phenotypically distinct populations of *Mtb in vivo*

Continuously changing environments within the tuberculous lung, the specific bacillary load, and a variable spectrum of lesions (3, 4, 16, 23, 29) argue that at least two distinct *Mtb* populations, live and dead cells, must exist *in vivo*. However, there is currently no method that distinguishes metabolically living from dead *Mtb* cells, although one study (9) reported that only living *Mtb* cells secrete MPT64. Here, we test the hypothesis that ZN staining combined with IHC for secreted *Mtb* antigens can identify phenotypically diverse *Mtb* populations *in vivo*. We examined testicular tissue whose seminiferous tubules contain numerous bacilli and are bordered by a basement membrane that separates the tubule from the sparsely infected surrounding tissue, which aids in evaluating the specificity of *Mtb* antigen immunostaining. H&E histology revealed the presence of substantial cellular debris (Figure E4A) which overlaps with *Mtb* RNAscope signals (Figure E4B, E3B). *Mtb* bacilli co-localize with strong Ag85B, ESAT-6, and uncharacterized cell surface protein (USP) positivity in sequential tissue slices (Figures E4C-E), suggesting that live bacilli produced these antigens. While these antigens are typically associated with the *Mtb* cell wall, clear delineation of the bacillary rod shape is not always possible. In some cases (*e.g.*, USP, ESAT-6), positivity was seen as scattered, brown patches, whereas some rod-like shapes (Figure E4D, E) were observed, consistent with previous studies (1, 11, 30, 31). These findings suggest that Ag85, ESAT-6, and USP staining patterns may not always match the rod-like morphology of *Mtb*.

To determine whether distinct microenvironments can contribute to differential antigen production, we examined whether all AFB produce Ag85B. Intriguingly, our dual ZN/Ag85B IHC stain demonstrated that not all AFB produce Ag85B. This was evident by numerous ZN-positive/Ag85B-negative bacilli close to Ag85B-positive bacilli (Figure 6A-D, E5A, B, E6). Ag85-positive and ZN-positive/Ag85B-negative bacilli were observed intracellularly and extracellularly within host cells that did, or did not, accumulate Ag85B (Figure 6E-I). Similar to Canetti (4), we also identified “ghost cells” that are weakly acid fast (Figure 6F).

**Figure 6.**
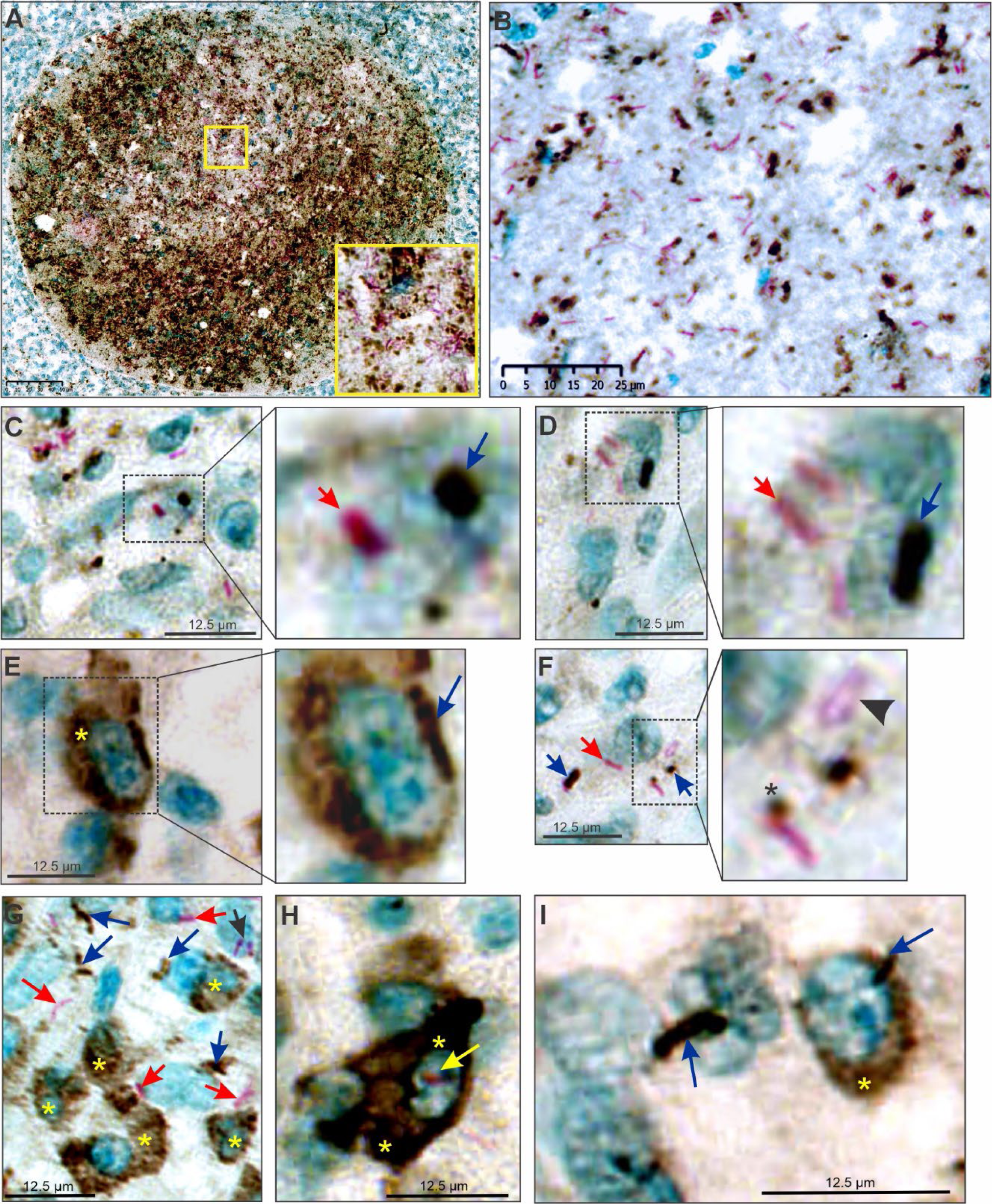
Phenotypically distinct *Mtb* populations exist within human extrapulmonary TB tissue. (**A**) Medium power image of a seminiferous tubule with combined ZN staining (pink) and Ag85B IHC staining (brown). Inset; note the presence of Ag85B-positive and -negative bacilli within the seminiferous tubule. (**B**) High power image of ZN/Ag85B staining showing Ag85B-positive bacilli near AFB-positive/Ag85B-negative bacilli. (**C, D**) High power images of ZN/Ag85B dual staining showing Ag85B-positive (blue arrow) and -negative (red arrow) bacilli. **(E)** Ag85B-positive *Mtb* bacillus (blue arrow) and cytoplasm (yellow asterisk). (**F**) weakly acid-fast ghost bacillus (arrowhead) and an AFB with Ag85B positivity at the one pole (asterisk). (**G**) intracellular and extracellular AFB (red arrows) and Ag85B-positive bacilli (blue arrows), some with Ag85B-positive host cell cytoplasm (yellow asterisks). (**H**) Ag85B-positive host cell cytoplasm (yellow asterisk) with Ag85B-negative *Mtb* bacillus (yellow arrow). (**I**) Two Ag85-positive *Mtb* bacilli (blue arrows); one is inside an Ag85B-positive host cell (yellow asterisk).

In sum, we demonstrate that two phenotypically distinct *Mtb* populations exist in human TB tissue: Ag85B-positive and Ag85-negative bacilli. This finding strongly suggests that Ag85B IHC alone may underestimate the number of bacilli. This has important implications for TB diagnosis and pathogenesis since mixed populations of *Mtb* may differently influence diagnosis, pathogenesis, and immunity.

### Using RNAscope and secreted *Mtb* antigens as tools for guiding TB therapy

To evaluate the potential of RNAscope to guide clinical intervention, we examined a 61-year-old female initially diagnosed with disseminated histoplasmosis. Biopsies of inguinal and retroperitoneal lymph nodes were performed 414 and 13 days before hospitalization, respectively (Figure 7A). Both biopsy specimens were ZN-negative and revealed granulomatous inflammation (Figure E7). On hospital day 18, bronchoscopy with bronchoalveolar lavage (BAL) showed a negative gram stain and AFB smear. The patient expired on hospital day 19 with the cause of death attributed to complications of sepsis. Autopsy revealed disseminated granulomatous inflammation of the lungs, peritoneum, liver, left kidney, bone marrow, spleen, and multiple central lymph nodes (Figure E8A-D). ZN staining showed diffuse involvement by AFB (Figure E9), and the cause of death was amended to be complications from disseminated TB. Culture of antemortem BAL fluid revealed the presence of pansensitive *Mtb* complex 47 days after hospital admission (Figure 7A).

**Figure 7.**
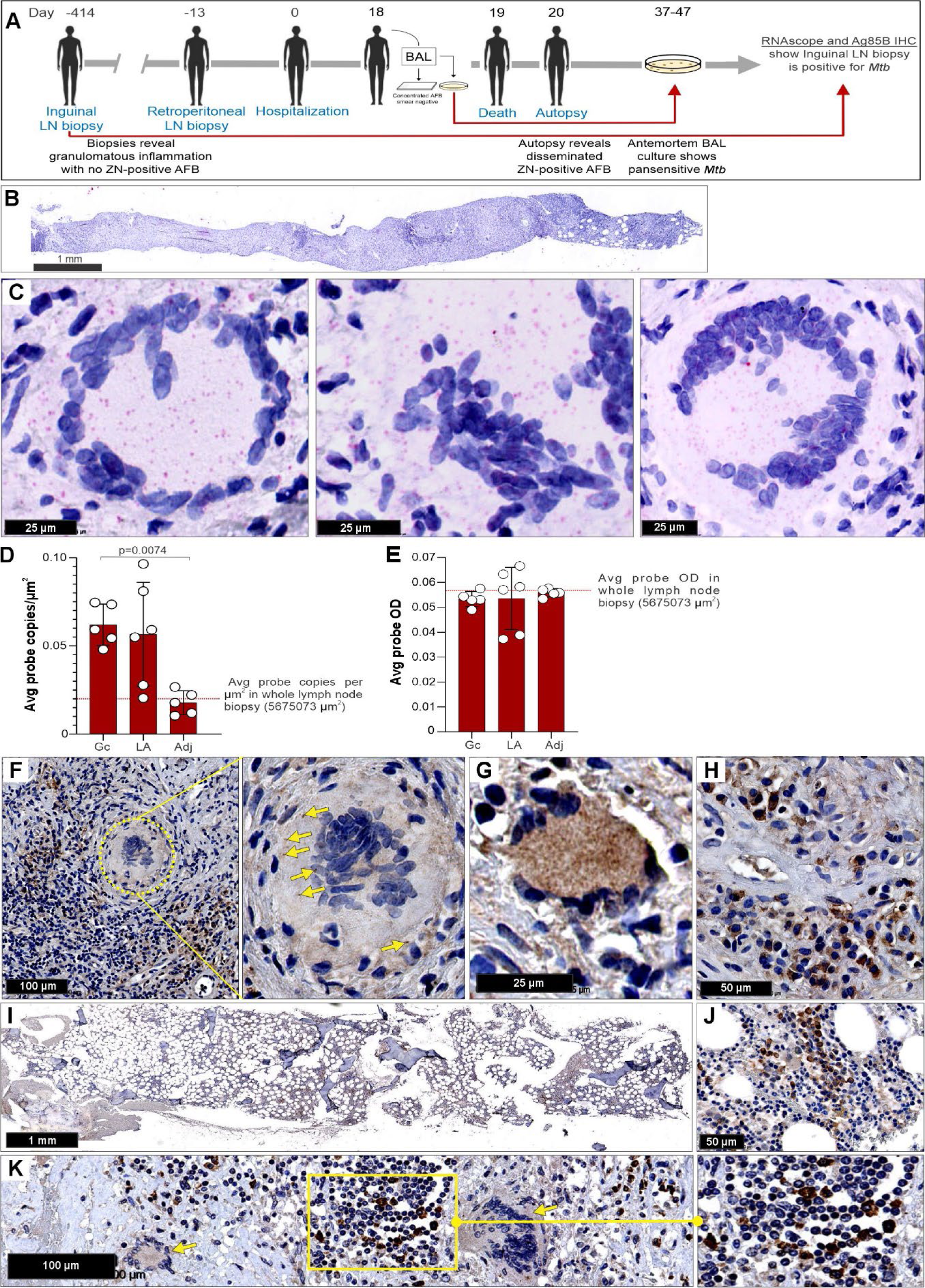
Application of RNAscope may help guide therapeutic intervention. (**A**) Flowchart depicting the biopsies, hospitalization, and postmortem analyses of a patient with undiagnosed TB. (**B**) Medium power image of an antemortem left inguinal lymph node needle biopsy specimen obtained 414 days prior to hospital admission. (**C**) High power images of the inguinal lymph node in (**B**) revealing *Mtb* RNAscope signals in and around giant cells. (**D**) Average number of RNAscope signals per square micrometer and (**E**) average probe optical density (OD) in and around giant cells (Gc), lymphocytic aggregates (LA) and adjacent (Adj) lymphoid tissue. (**F**) Ag85B-positive staining in and around giant cells in the inguinal lymph node specimen. The circled area shows weakly positive giant cells with several Ag85B-positive bacilli (yellow arrows). (**G**) A strongly Ag85B-positive giant cell in the inguinal lymph node specimen. (**H**) High power image of Ag85B accumulation within the cytoplasm of lymphocytes in the inguinal lymph node specimen. (**I**) Low power image of Ag85B-positive cells in the antemortem bone marrow biopsy. (**J**) Medium power image showing Ag85B positivity in the cytoplasm of lymphocytes within bone marrow. (**K**) Ag85B positivity in the postmortem periaortic lymph node specimen. Notably, specimens in **B, C and F-J** are ZN-negative. Data in (**D**) and (**E**) represent the mean ± SD and each data point represents the number of RNAscope signals per square micrometer (**D**) or probe OD (**E**) in each zone (Gc, LA, and Adj) of the inguinal lymph node (n=5-6). Data in (**D**) were analyzed using one-way ANOVA and Bonferroni’s multiple comparison test.

With longitudinal specimens available, we applied RNAscope to the ZN-negative inguinal lymph node biopsy obtained 414 days prior to hospital admission (Figure 7B) and found numerous *Mtb* mRNA transcripts (Figure 7C). Quantitation using HALO® analysis showed a higher APC around and inside giant cells and lymphocytic aggregates compared to adjacent lymphoid tissue and the whole biopsy (Figure 7D) with no difference in average probe optical density (OD) (Figure 7E). This suggests that the *Mtb* APC can help predict pathophysiological abnormalities in human TB tissue.

The detection of *Mtb* via RNAscope is further supported by strong Ag85B positivity in giant cells (Figure 7F, G) and surrounding lymphocytes (Figure 7F) in the antemortem inguinal lymph node specimen. Antemortem bone marrow biopsy (Figure 7J) and postmortem periaortic lymph node (Figure 7K) specimens also demonstrated Ag85B positivity.

In this clinical scenario, application of RNAscope detection of *Mtb* mRNA and Ag85 IHC to biopsy tissue could have positively identified *Mtb*, providing much earlier diagnosis of disseminated TB, possibly enabling effective TB treatment. These findings suggest that detection of *Mtb* mRNA and antigens has considerable diagnostic value.

## DISCUSSION

Major unmet needs in the TB field are the ability to consistently identify *Mtb* bacilli and the means to gain a clear understanding of how *Mtb* contributes to human tissue pathology. We have taken initial steps toward addressing these needs by adapting the RNAscope platform to identify intact and disintegrating *Mtb* bacilli and single *Mtb* transcripts in human TB tissues. We show that large quantities of *Mtb* mRNA are present in ZN-negative lung tissue specimens from a confirmed pulmonary TB case, that *Mtb* mRNA is found intracellularly and extracellularly, and that *Mtb* mRNA and antigens accumulate in cells from histologically normal and abnormal tissue. We also provide evidence of two phenotypically distinct *Mtb* cell populations *in vivo*. Lastly, our antemortem biopsy studies provide strong evidence that RNAscope can help guide therapeutic intervention. Overall, the ability of RNAscope to detect bacilli in diverse morphological states and monitor its historical footprints *in vivo* advances our understanding of TB pathophysiology and diagnosis.

RNAscope has diagnostic potential due to its highly sensitive and specific probe design. In particular, the detection of clearly-defined bacillary rod shapes that are visually and morphologically similar to positive ZN staining as well as punctate dots indicating single transcripts makes it an attractive diagnostic platform. Current dogma asserts that bacterial mRNA is an unlikely candidate due to its susceptibility to degradation, which raises the question of *in vivo* mRNA stability outside of the bacillus, *i.e*., inside or outside the host cell. Stability of *Mtb* mRNA is likely influenced by the specific host organelle (*e.g*., phagosome) in which it is contained. Bacteria contain multienzyme RNA degradomes comprised of endo- and exoribonucleases, RNA helicases, and metabolic enzymes that degrade RNA during growth and in response to environmental cues (32, 33). While the degradome is highly effective in degrading intra-bacterial mRNA, very little is known about the stability of extracellular bacterial mRNA within and outside the infected host cell. As RNAscope is effective in detecting extracellular *Mtb* mRNA, our findings suggest that mRNA released from the bacterial cell is sufficiently stable to be detected by this platform.

Pathologists are frequently confronted with diagnostic challenges in TB cases since most granulomas contain few, if any bacilli. Since RNAscope detected *Mtb* mRNA in AFB-negative lung and lymph node specimens from confirmed TB patients, this technique could help confirm diagnosis by a pathologist to guide the treating clinician. Furthermore, the interventional value of RNAscope was demonstrated by confirming the presence of *Mtb* mRNA in a ZN-negative antemortem biopsy from a patient initially diagnosed with histoplasmosis. Hence, RNAscope analysis could have guided potentially life-saving therapy.

A longstanding gap in our understanding of TB is whether *Mtb* exists in more than one phenotypic state *in vivo*, and if so, how these phenotypes could be detected. It is logical to assume that *Mtb* exists as live and dead bacilli in human tissue (3); however, this is difficult to demonstrate experimentally and no successful method has been reported to date. *Mtb* naturally exists in fully, partially and non-acid fast states and loses its acid fastness during disintegration (4, 5). Hence, acid fastness is not an indicator of viability. Several studies have shown that *Mtb* surface and/or secreted antigens can be detected in human sputum (9) and in human (1, 11, 31, 34, 35) and guinea pig (6) tissue specimens. We identified two distinct populations of *Mtb* in human specimens: those that produce Ag85B and those which do not. Since many of these bacilli are in close proximity, differential *ag85B* expression in response to different environmental signals is unlikely. Since Ag85B secretion is not universal, Ag85B IHC alone is likely to underestimate bacillary burden, consistent with prior studies (30).

An unexpected discovery was the accumulation of *Mtb* mRNA in the cytoplasm of HAMR cells. This important finding is reminiscent of *in situ* hybridization studies that demonstrated the presence of *Mtb* DNA in host cells (19, 36). We were unable to identify discernable bacilli inside these cells, suggesting that the source of mRNA is disintegrated intracellular bacilli. An alternative explanation is that HAMR cells take up exogenous *Mtb* RNA and/or secreted antigens. Recent studies have shown that *Mtb* can produce extracellular vesicles (EVs) that contain hundreds of proteins, including Ag85A, (37) secrete RNA *in vitro* (38) and release mRNA into the cytosol in macrophages (28). Further, EVs from *Mtb*-infected macrophages contain *Mtb* mRNA (28, 39). Hence, uptake of *Mtb*-derived EVs by HAMR cells could account for the accumulation of RNA and Ag85B. RNA from *Streptococcus agalactiae* (group B streptococcus) has been implicated as an immunomodulatory PAMP that induces IFN-β production in dendritic cells (25), suggesting that *Mtb* nucleic acids and/or secreted antigens may also exert similar effects. Indeed, *Mtb* RNA is sensed in a manner dependent on melanoma differentiation factor 5 (MDA-5), an RNA sensor in the RIG-I-like 3 receptor family, resulting in increased IL-1β production, inflammasome activation and attenuation of autophagy. Notably, these effects culminate in increased *Mtb* survival in macrophages (27). Overall, these studies suggest mechanisms by which HAMR cells in TB patients may accumulate *Mtb* mRNA and/or secreted antigens and provide context for investigating of the fate and immunomodulatory function of these cells.

Our finding of *Mtb* mRNA in bronchial epithelial cells provides new insight into *Mtb* tropism, as this microenvironment is not routinely examined by pathologists. The accumulation of *Mtb* mRNA inside bronchial epithelium indicates significant bacillary destruction in this environment, which was confirmed by Ag85B positivity, and is consistent with our finding that *Mtb* can cross the bronchial epithelium (29).

We have demonstrated the potential of RNAscope to detect *Mtb* mRNA within intact bacilli, disintegrating bacilli and single *Mtb* transcripts in a range of human tissues including pulmonary, lymphatic and testicular specimens. Our case study involving ante- and postmortem biopsy specimens provides compelling evidence that RNAscope has potential to reduce clinical time-to-diagnosis. Positive Ag85B immunostaining confirmed the RNAscope results, increasing diagnostic confidence. In cases where clinical suspicion remains high despite negative AFB staining, RNAscope has the potential to confirm or exclude the diagnosis of TB.

Our study has limitations that could affect its application. Firstly, distinguishing viable from nonviable bacilli remains to be demonstrated since the presence of mRNA is not necessarily an indication of viability. Secondly, the design and synthesis of RNAscope probes is relatively expensive, which may limit its initial clinical application.

In conclusion, our findings have important implications for TB pathophysiology and diagnosis. RNAscope detected *Mtb* mRNA *in vivo*, which was previously thought to be unstable, as well as in intact and disintegrating bacilli. We also detected *Mtb* RNA within ZN-negative human tissues obtained from pulmonary and extrapulmonary sites and found that *Mtb* mRNA accumulates within some host cells but is also present outside host cells. Additionally, we show that antigen secretion is not universal, demonstrating two diverse populations of *Mtb in vivo*. These findings imply applications that could have considerable impact, such as the early identification of individuals who are at risk of dying from TB-related causes and tracking the clinical course of disease via the historical footprints of *Mtb*.

