## Supplementary material for "Detection of *Mycobacterium tuberculosis* in human tissue via RNA *in situ* hybridization": Online Data Supplement

### Detection of *Mycobacterium tuberculosis* in human pulmonary and extrapulmonary tissue via RNA *in situ* hybridization

#### SUPPLEMENTAL MATERIALS AND METHODS

##### Ethics and Human Subjects

This study was approved by the University of KwaZulu-Natal Biomedical Research Ethics Committee (BREC; class approval study number BCA 535/16, and BE019/13) and the University of Alabama Birmingham Institutional Review Board (IRB; study numbers IRB-300008174 and IRB-300008174-2). Consent for research use of the autopsy material is included in the UAB authorization for autopsy consent form signed by the decedent's next of kin. Patients undergoing lung resection for TB were recruited from King DinuZulu Hospital Complex, a tertiary center for patients with TB in Durban, South Africa. *Mtb* infected human lung tissues are routinely obtained after surgery for removal of irreversibly damaged lobes or lungs (bronchiectasis and/or cavitary lung disease). Written informed consent was obtained from all participants. All patients undergoing lung resection for TB had completed a full 6- to 9- month course of anti-TB treatment or up to two years of treatment for drug-resistant TB. Patients were assessed for the extent of pulmonary disease (cavitation and/or bronchiectasis) via HRCT. The fitness of each patient to withstand a thoracotomy and lung resection was determined by using the Karnofsky score, 6-minute-walk test, spirometry, and arterial blood gas measurement. Assessment of patients with massive hemoptysis included their general condition, effort tolerance before hemoptysis, arterial blood gas measurement, serum albumin concentration, and HRCT imaging of the chest. On gross assessment, all pneumonectomies or lobectomies were bronchiectatic, hemorrhagic, variably fibrotic, and atelectatic and contained visible tubercles. See Table E1 in the online data supplement for description of human subjects and tissues.

##### Histology

Human tissue specimens (pulmonary, testicular, neonatal) were aseptically removed and fixed in 10% neutral buffered formalin (10% NBF). Specimens were processed in a vacuum filtration tissue processor using a xylene-free protocol. Tissue sections were embedded and blocked-in paraffin wax.

Sections were cut at 4 μm, baked at 56 °C for 15 min, dewaxed through 2 changes of xylene and rehydrated through descending grades of alcohol to water. Routine H&E staining was performed by placing slides in hematoxylin for 5 min, washing in tap water for 2 min, bluing in lithium carbonate for 1 min, rinsing in tap water for 2 min and counter staining with eosin for 5 min before a final rinse in tap water for 2 min. Slides were dehydrated in ascending grades of alcohol, cleared in xylene, and coverslip mounted with DPX (Distyrene, Plasticizer, and Xylene).

**Immunohistochemistry**

Human lung tissue was cut into 2 μm thick sections, mounted on charged slides, and baked at 56 °C for 15 min on a hotplate. Mounted sections were dewaxed in 2 changes of xylene followed by rinsing in 2 changes of 100% ethanol and 1 change of SVR (95%). Slides were then rinsed in tap water for 2 min followed by antigen retrieval via Heat Induced Epitope Retrieval (HIER) in Tri-Sodium Citrate (pH 6.0) for 30 min. Slides were cooled for 15 min and rinsed in tap water for 2 min. Endogenous peroxide activity was blocked using 3% hydrogen peroxide (Novolink) for 10 min at room temperature (RT). Slides were then rinsed in PBST and blocked with protein block (Novolink) for 5 min at RT. Sections were incubated with primary antibody directed against *Mtb* antigen 85B (Ag85B; Abcam cat # ab43019,1:500), *Mtb* Early Secreted Antigenic Target 6 (ESAT-6; Abcam cat # ab26246, 1:500) or *Mtb* Uncharacterized Surface Protein (USP; Lifespan Bioscience cat # LS-C 683286, 1:500) followed by rinsing in PBST and incubated with the polymer (Novolink) for 30 min at RT. Slides were then rinsed and stained with Diaminobenzidine (DAB) for 5 min, rinsed under running water and counterstained with hematoxylin for 2 min. Slides were rinsed in tap water, blued in 3% ammoniated water for 30 s, rinsed in tap water, dehydrated in ascending grades of alcohol, cleared in xylene, and coverslip mounted with DPX (Distyrene, Plasticizer, and Xylene).

**Microscopy and Imaging**

Microscope slides stained with H&E, ZN, IHC (*Mtb* Ag85B, ESAT-6, USP) or red RNAScope chromogen were imaged using a Hamamatsu NDP slide scanner (NanoZoomer RS2, Model C10730-12) using its viewing platform (NDP.View2). Imaging of clinical H&E, ZN, and GMS tissue staining was performed using an Olympus BX50 microscope with a DP23 digital microscope camera using Olympus cellSens Entry 3.2 image acquisition software (Build 23706). Automatic exposure settings were used after manual white balance. Image resolution was set at “Full 3088 x 2076”.

**RNAscope**

The RNAscope 2.5 High Definition (HD)-Red assay kit (Advanced Cell Diagnostics (ACD) cat # 322350) employed here uses a red chromogen to visualize target mRNA. All RNAscope probe sets were purchased from ACD. To ensure probe set specificity, a probe design algorithm is used examine each probe for potential cross-reactivity against the host cell’s transcriptome (for detecting single mRNAs within eukaryotic or prokaryotic cells) or against transcriptomes of related organisms (for identifying specific organisms). We used a positive control RNAscope probe set of 16 probe pairs directed against bp 139-989 of human peptidylprolyl isomerase B (*PPIB*) mRNA (ACD cat # 313901). The negative control probe set consists of 10 probe pairs directed against bp 414 -862 of *Bacillus subtilis* dihydrodipicolinate reductase (*dapB*) mRNA (ACD cat # 310043). The *Mtb*-specific probe set (ACD cat # 552911) consists of 120 probe pairs directed against mRNAs from six genes (20 probe pairs per gene): secreted L-alanine dehydrogenase (*ald*), catalase-peroxidase-peroxynitritase T (*katG*), trehalose-6-phosphate phosphatase (*otsB1*), resuscitation-promoting factor A (*rpfA*), resuscitation-promoting factor B (*rpfB*), and ATP-binding protein (ABC transporter) (*irtB*).

Human lung, testicle, lymph node and neonatal tissue blocks were cut at 4 µm and mounted on charged slides. The sections were then deparaffinized in RNAscope deparaffinization solution following the manufacturer's protocol. Slides were then rehydrated rinsing twice in 100% ethanol for 5 minutes each, followed by two rinses in 70 % ethanol for 5 minutes each, and two rinses in distilled water for 5 minutes each. Slides were subjected to antigen retrieval by incubating the tissue sections in RNAscope Target Retrieval Reagent at 95-99°C for 15 minutes for our control tissue and 20 minutes for test lung tissue. Both test and control slides underwent protease treatment by applying RNAscope Protease Plus solution onto each tissue section at 40 °C for 30 minutes to permeabilize the cells and increase probe accessibility. This was followed by probe hybridization was applied to the tissue sections to hybridize to the target mRNA according to the manufacturer's protocol. Slides were then subjected to a series of signal amplification steps using an RNAscope 2.5 HD-Red assay kit according to the manufacturer's protocol. Tissue sections were then counterstained with hematoxylin to visualize cell nuclei. Slides were mounted using RNAscope Vector mount medium that is compatible with the red chromogen used in the assay. Imaging was performed on a slide scanner as stated above.

**HALO® analysis**

Whole-slide image analysis was performed using Halogen-Assisted Light Optimization (HALO) v3.6.4134 (Indica Labs, Corrales, NM). HALO® employs advanced image analysis algorithms to automate the analysis of RNAscope signals in TB tissue specimens. It is equipped with advanced quantification features, such as calculating the number of RNA molecules, determining their intensity, and assessing their spatial distribution. Regions of interest (ROIs) were drawn on the tissue using the annotation tools provided in the HALO® platform. The ISH module v4.2.3 (Indica Labs, Corrales, NM) was used to detect the RNAscope probe signals. For visualization, the Area Quantification module v2.4 was used. For both modules, the magnification was set to "1" with the parameters described in Table E2.

**Statistics**

Results are shown as Mean ± SD. All graphs were plotted using GraphPad Prism v6.04 (GraphPad Software Inc., USA) and the statistical significance was calculated by applying Student's t-test and One-way ANOVA. Significance was accepted at a *P* value less than 0.05. Exact *P* values have been included in the data plots for statistical evaluation.

#### SUPPLEMENTAL FIGURES

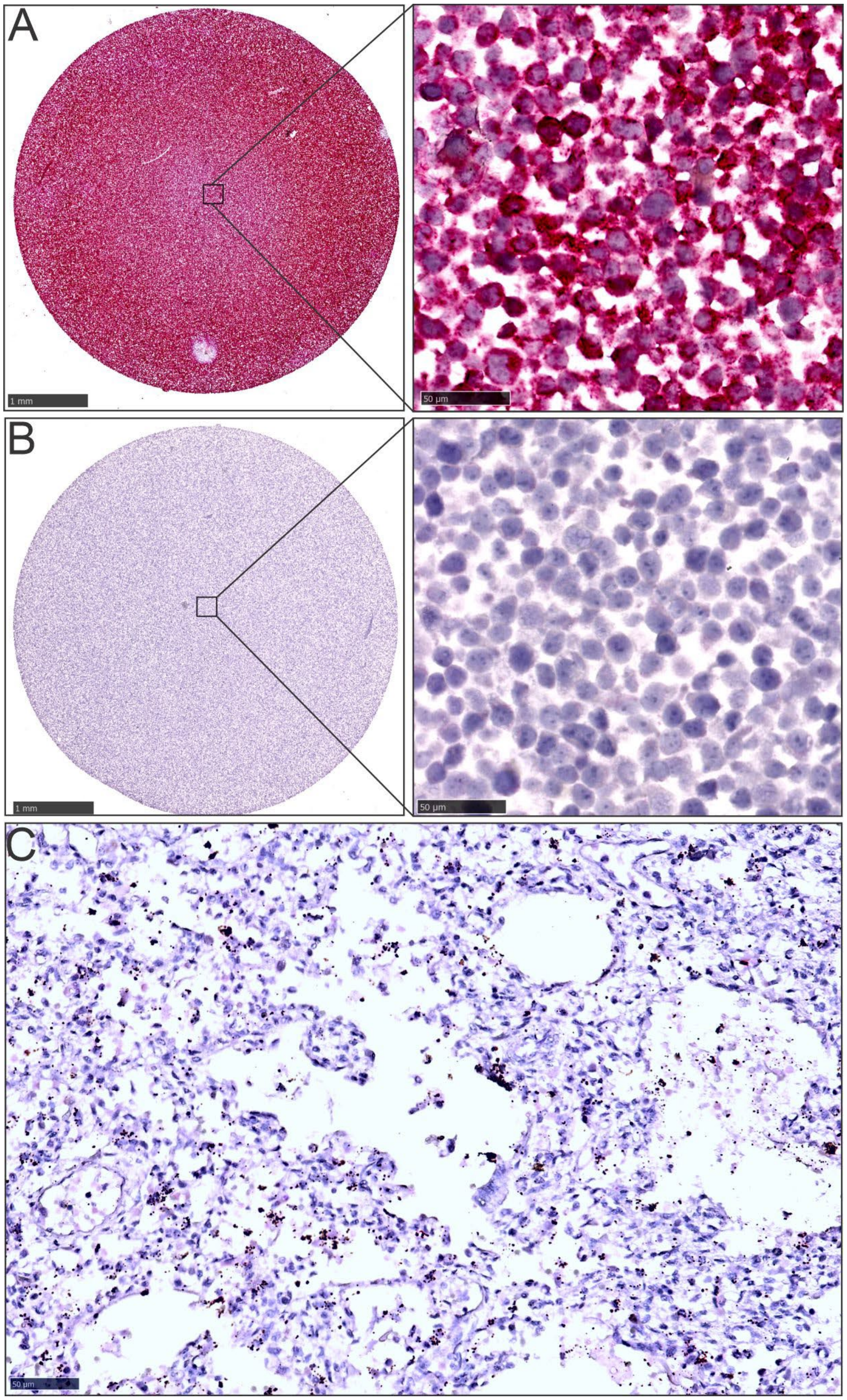

**Figure E1. RNAscope probe set and tissue control assays.** (**A**) Hela cell pellet thin section exposed to a positive control probe set directed towards human peptidylprolyl isomerase B (*PPIB*) mRNA show a strong positive signal. (**B**) Hela cell pellet section exposed to a negative control probe set specific for *Bacillus subtilis* dihydrodipicolinate reductase (*dapB*) mRNA is RNAscope signal negative. (**C**) Human neonatal lung tissue exposed to the *Mtb*-specific probe set is RNAscope signal negative.

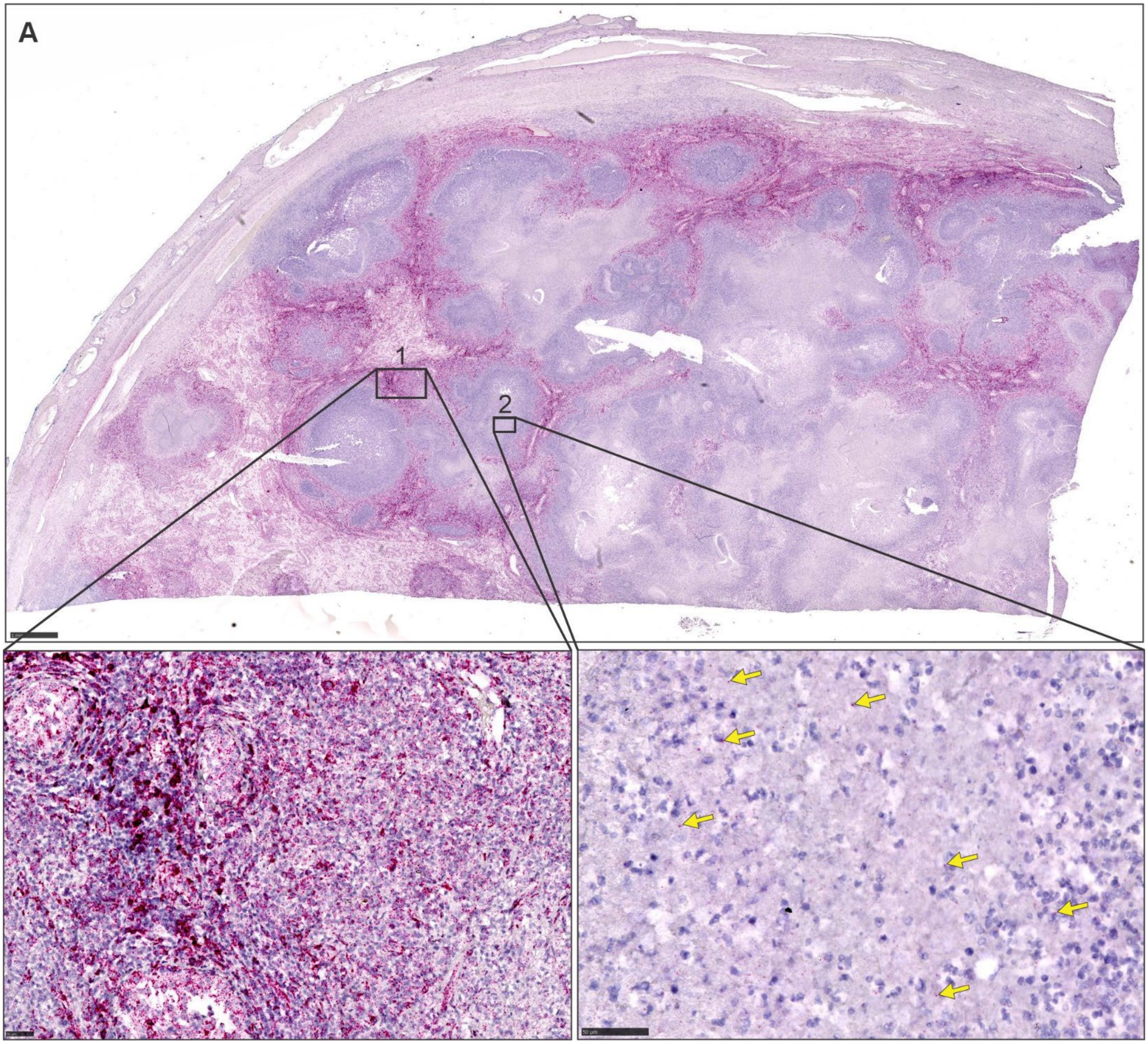

**Figure E2. RNAscope tissue and probe set control assays.** (**A**) Low power image of RNAscope signals from a human mRNA positive control probe set directed against peptidylprolyl isomerase B (*PPIB*) mRNA in a testicular specimen from a TB patient. Insets; medium power images of boxed areas in (**A**). (1) Medium power image showing an abundance of RNAscope positive (*PPIB*) signals. (2) Sparse RNAscope signals in the necrotic area are indicated by yellow arrows.

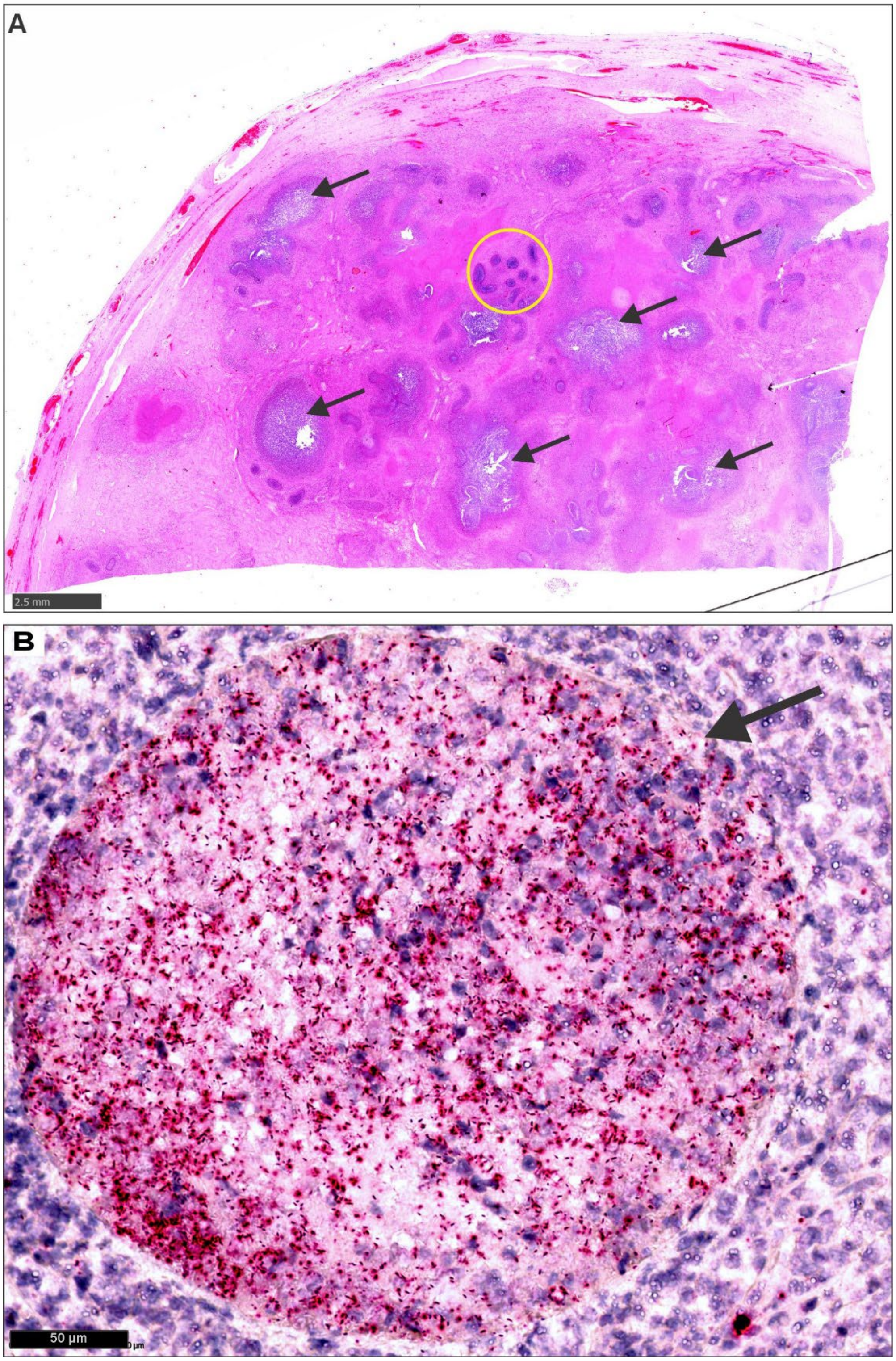

**Figure E3. RNAscope detects *Mtb* in seminiferous tubules.** (**A**) Low power image of an H&E-stained testicular specimen obtained from a TB patient. Circled area; seminiferous tubules. Arrows indicate granuloma. (**B**) Medium power image of a seminiferous tubule containing numerous *Mtb*-specific *Mtb* RNAscope signals. Black arrow indicates the tubule basement membrane.

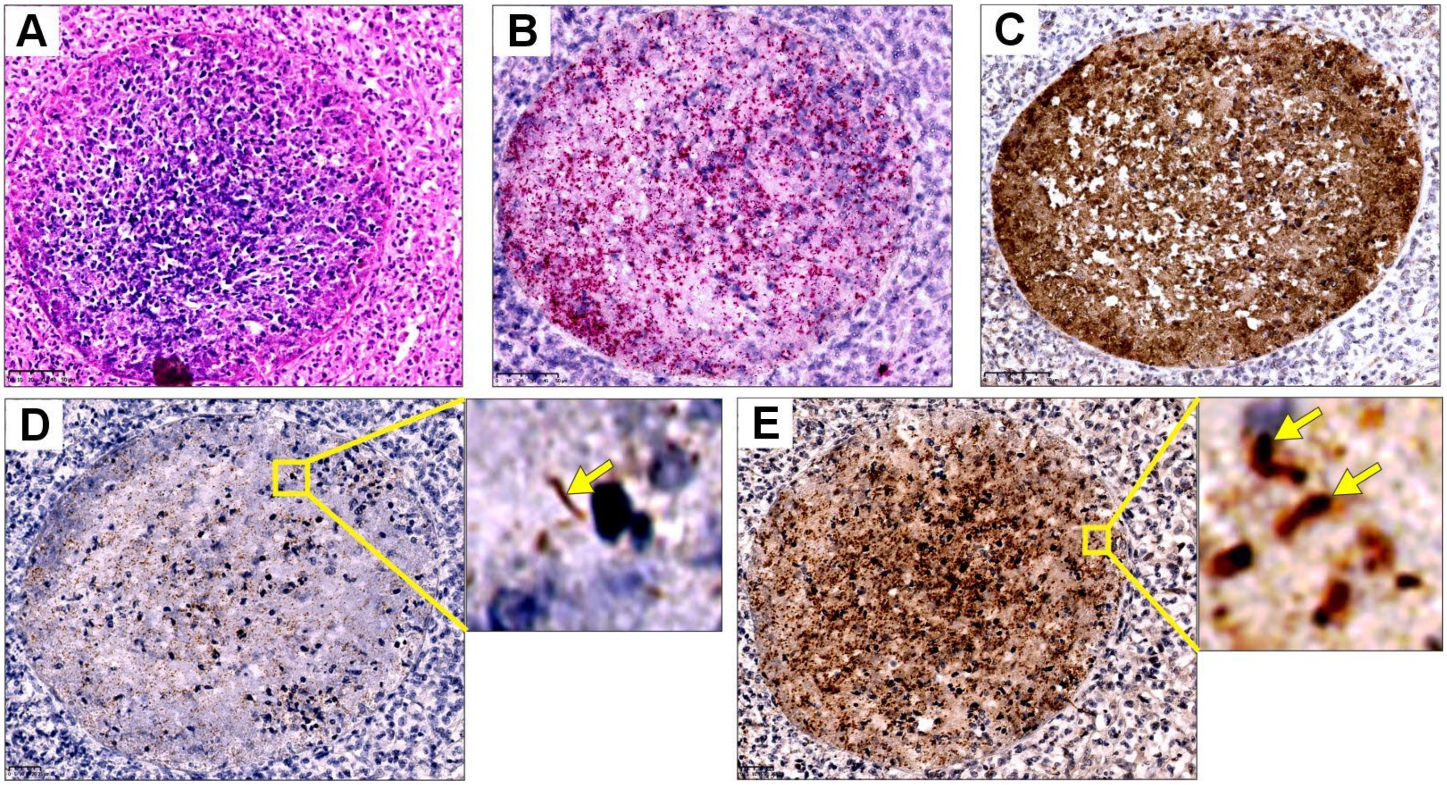

**Figure E4. Accumulation of *Mtb* antigens in extrapulmonary TB tissue.** Images of consecutive sections of a seminiferous tubule from a TB patient. (**A**) Medium power image of H&E staining. (**B**) Medium power image showing *Mtb*-specific RNAscope signals (same image as Figure E3B). (**C**) Medium power image of *Mtb* USP-positive IHC staining. (**D**) Medium power image of ESAT-6-positive IHC staining with a high-power image depicting ESAT-6-positive bacillary shapes (yellow arrow). (**E**) Medium power image of Ag85B-positive IHC staining with high power image of Ag85B-positive bacillary shapes (yellow arrows). (**C**, **D**, **E**) Note the clear line of demarcation formed by the tubule basement membrane between the accumulated secreted antigens inside the seminiferous tubule and the surrounding tissue confirming the specificity of the antibodies.

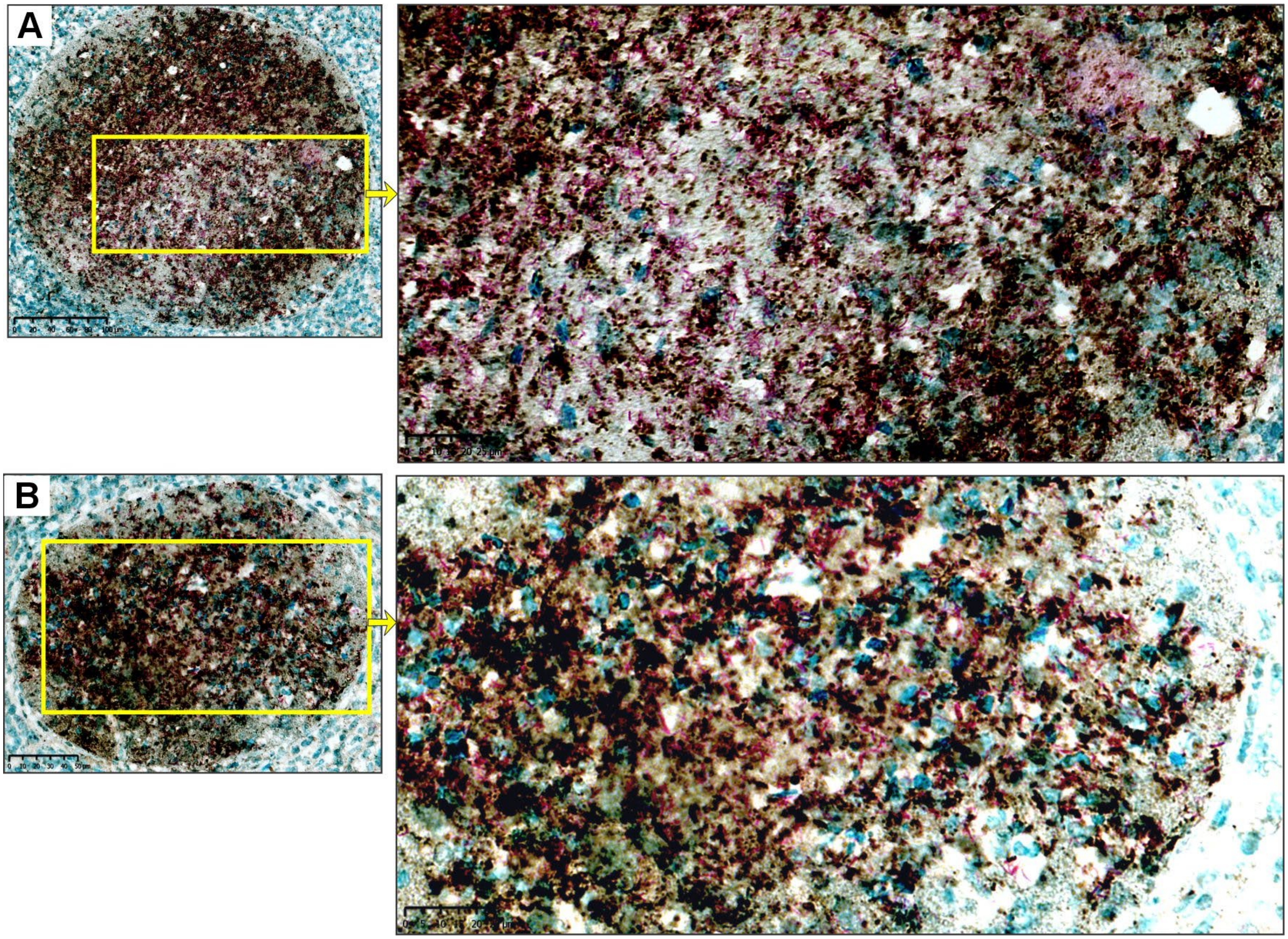

**Figure E5. *Mtb* Ag85B-positive and -negative bacilli inside seminiferous tubules.** (**A**) and (**B**) low power images of seminiferous tubules from a TB patient with combined ZN staining (pink) and Ag85B IHC staining (brown). Insets; medium power images illustrating Ag85B-positive and ZN-positive/Ag85B-negative bacilli.

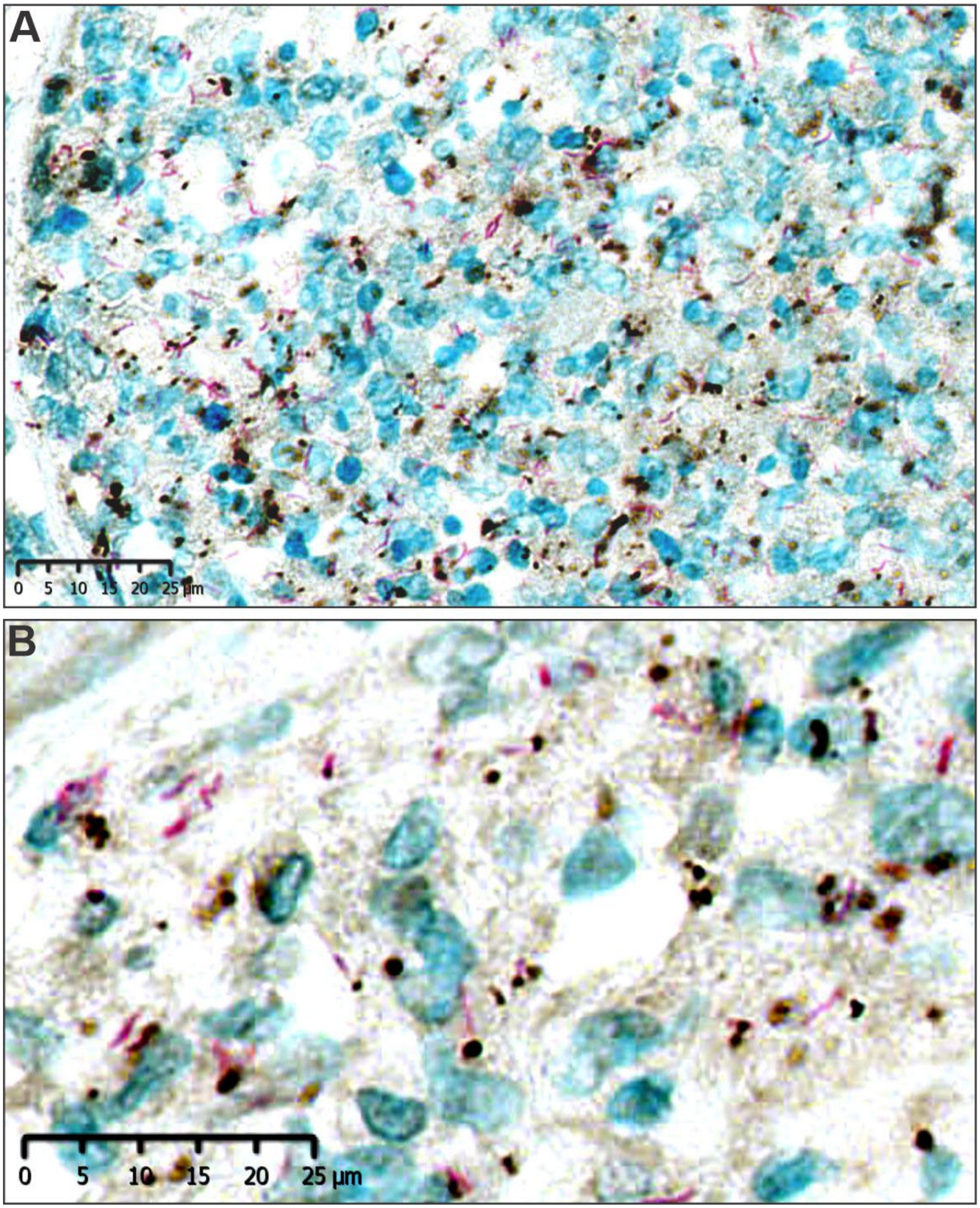

**Figure E6. Ag85B-positive and -negative *Mtb* bacilli in extrapulmonary TB tissue.** (**A**) Low power and (**B**) medium power images of testicular tissue (outside the seminiferous vesicle) with combined ZN staining (pink) and Ag85B IHC staining (brown).

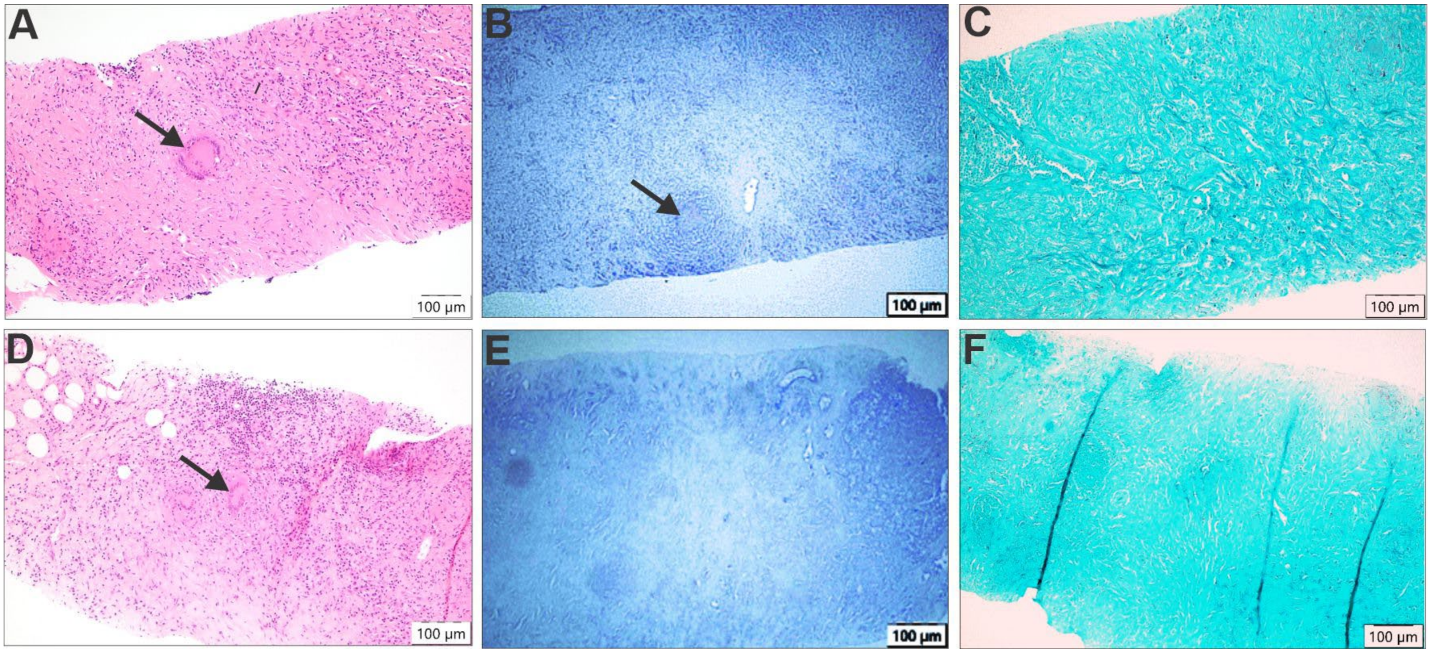

**Figure E7. Human antemortem biopsy specimens showing evidence of granulomatous inflammation but absence of AFB and fungal organisms.** (**A**-**C**) Left inguinal lymph node biopsy collected 414 days prior to hospital admission. (**D**-**F**) Retroperitoneal lymph node biopsy collected 13 days prior to hospital admission. (**A** and **D**) H&E staining. (**B** and **E**) ZN staining shows no evidence of AFB. (**C** and **F**) Gomori Methenamine Silver (GMS) staining shows no evidence of fungus or yeast. Black arrows indicate giant cells associated with granuloma.

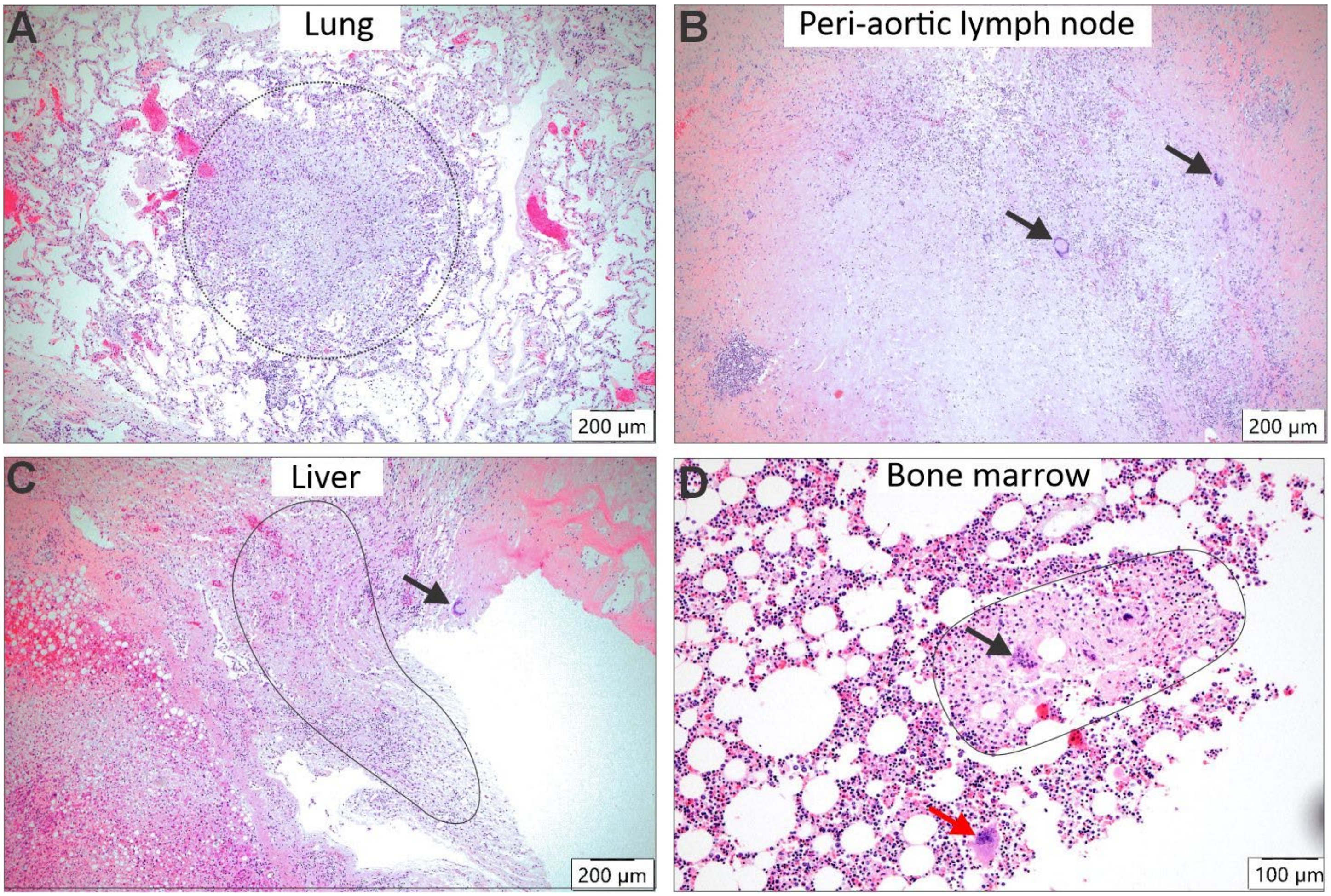

**Figure E8. Diffuse granulomatous inflammation in multiple tissues collected at autopsy**. H&E staining of (**A**) Active granuloma with necrosis in the lung (circled). (**B**) Peri-aortic lymph node with giant cells (black arrows). (**C**) Liver specimen with necrotic fibrin exudate (circled) and giant cell (black arrow). (**D**) Organized granuloma (circled) in bone marrow with giant cell (black arrow). Red arrow, megakaryocyte.

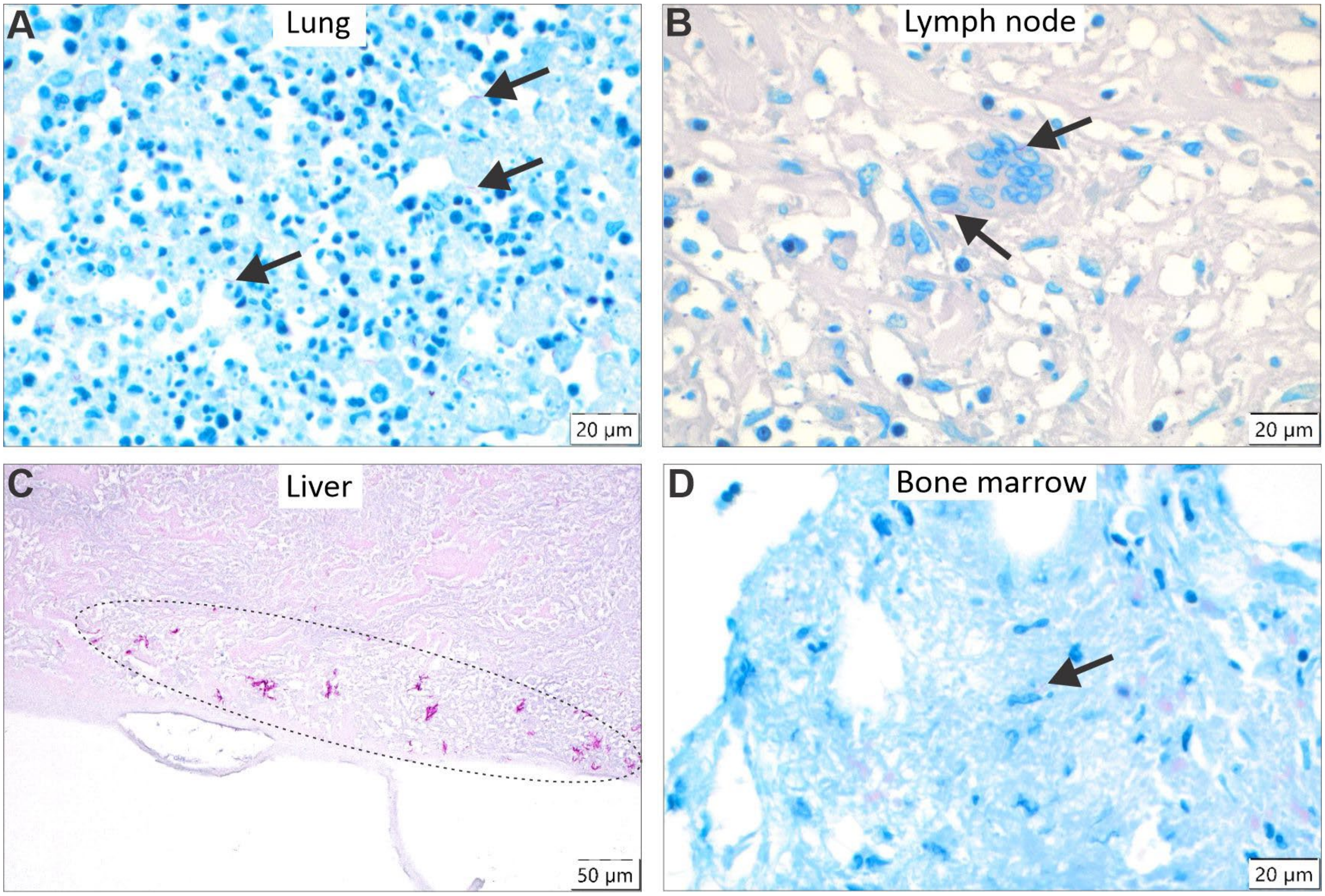

**Figure E9. Acid fast bacilli in multiple tissues collected at autopsy.** ZN staining of autopsy specimens from (**A**) Lung, (**B**) Peri-aortic abdominal lymph node, (**C**) Liver, and (**D**) Bone marrow. Black arrows and area within oval indicate ZN-positive *Mtb* bacilli. ZN stain includes methylene blue counter stain. Liver section displayed surface-localized necrotic fibrin exudate that prevented nuclear staining.

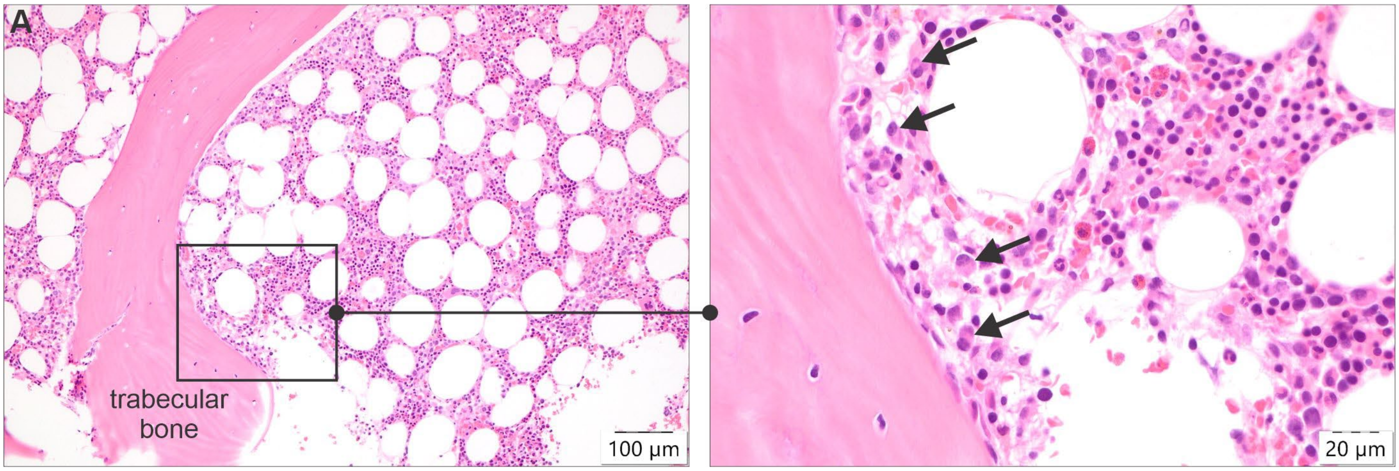

**Figure E10. Bone marrow biopsy on hospital day four.** (**A**) H&E at low (10x) magnification. Inset, medium (40x) magnification. Black arrows indicate plasma cells.

**Table E1. Clinical characteristics of human subjects.**

| # | Patient # | Age | Sex | Tissue specimen | Macroscopic and microscopic features |
|---|---|---|---|---|---|
| 1 | SL314-23 | 67 | M | Lobectomy, left upper lobe | Lobe weighing 323 g, pleural surface was hemorrhagic with patchy areas of caseous necrosis. Pathology features are those of necrotizing granulomatous inflammation with AFB, consistent with active TB. |
| 2 | SL463-23 | 38 | F | Lobectomy, left lung | Lobe weighing 281 g, cut sections showed evidence of bronchiectasis and caseative necrosis, a cavity was noted. Areas of necrotizing granulomatous inflammation were evident, and AFB were identified. Suppurative granulomas with microabscess formation were noted. |
| 4 | SL114-23 | 30 | F | Pneumonectomy, left Lung | Left lung weighing 229 g, lung was shrunken, fibrotic and cavitated with miliary tuberculosis. Numerous AFB were present, features of lymphoid interstitial pneumonia were present. |
| 5 | SL038-23 | 38 | M | Left orchidectomy | Left testis weighing 108 g, upon cut section, architectural distortion by extensive necrotizing granulomatous inflammation was demonstrated. Granulomas showed central caseative necrosis, blood vessels demonstrate extensive medial hypertrophy with luminal occlusion. Numerous AFBs were identified. Features are those of tuberculous epididymo-orchitis. |
| 6 | SLPM1-23 | | F | Neonatal lung tissue | Early neonatal death shortly after birth and prior to administration of BCG vaccination. Routine post-mortem lung samples (minimum 1 section from each lobe of the right and left lungs). Normal anatomy with no gross pulmonary pathology. Microscopy revealed variable alveolar expansion with patchy intra-alveolar oedema and hemorrhage. Infective pathogens, granulomata or neoplastic infiltrates were not seen. |
| 7 | 21-106 | 61 | F | Ante-mortem: Inguinal lymph node, retroperitoneal lymph node, bone marrow.<br><br>Post-mortem:<br>Lungs, peri-aortic lymph node, liver, and bone marrow. | Ante-mortem lymph nodes biopsies revealed granulomatous inflammation on H&E and were negative for AFB on ZN. Antemortem bone marrow revealed focal paratrabecular plasma cells and was not ZN stained clinically due to no granulomas present.<br><br>Post-mortem lungs tissue, periaortic lymph node, liver, and bone marrow all revealed diffuse granulomatous inflammation and were positive for AFB on ZN staining. |

**Table E2. Parameters and values for RNAscope signal quantitation using the HALO software platform**

| Parameters | Values |
|---|---|
| **Stain Selection** | |
| Number of Probes | 1 |
| Set the nuclear stain | 0.644, 0.716, 0.267 |
| Probe Name | RNA Probe |
| Set the RNA Probe stain | 0.242, 1.151, 0.452 |
| Set the exclusion stain | 2.407, 2.407, 2.407 |
| Output Image | RNA Probe Markup |
| **Optimize spot/signal detection** | |
| RNA Probe Markup Color | 255, 0, 0 |
| RNA Probe Contrast Threshold | 0.02 |
| RNA Probe Minimum Optical Density | 0.053 |
| Spot Segmentation Aggressiveness | 0.95 |
| RNA Probe Spot Size | 0.8, 20 |
| RNA Probe Size | 2.1 or 1.8 |
| Fill RNA Probe Holes | True |
| Output Image | RNA Probe Markup |
| **Set Exclusion Marker** | |
| Exclusion Threshold | 0.2 |
| Exclusion Radius | 0.5 |
